# Coisolation of peptide pairs for peptide identification and MS/MS-based quantification

**DOI:** 10.1101/2022.04.18.488679

**Authors:** Ian R. Smith, Jimmy K. Eng, Anthony S. Barente, Alexander Hogrebe, Ariadna Llovet, Ricard A. Rodriguez-Mias, Judit Villén

## Abstract

SILAC-based metabolic labeling is a widely adopted proteomics approach that enables quantitative comparisons among a variety of experimental conditions. Despite its quantitative capacity, SILAC experiments analyzed with data dependent acquisition (DDA) do not fully leverage peptide pair information for identification and suffer from undersampling compared to label-free proteomic experiments. Herein, we developed a data dependent acquisition strategy that coisolates and fragments SILAC peptide pairs and uses y-ions for their relative quantification. To facilitate the analysis of this type of data, we adapted the Comet sequence database search engine to make use of SILAC peptide paired fragments and developed a tool to annotate and quantify MS/MS spectra of coisolated SILAC pairs. In an initial feasibility experiment, this peptide pair coisolation approach generally improved expectation scores compared to the traditional DDA approach. Fragment ion quantification performed similarly well to precursor quantification in the MS1 and achieved more quantifications. Lastly, our method enables reliable MS/MS quantification of SILAC proteome mixtures with overlapping isotopic distributions, which are difficult to deconvolute in MS1-based quantification. This study demonstrates the initial feasibility of the coisolation approach. Coupling this approach with intelligent acquisition strategies has the potential to improve SILAC peptide sampling and quantification.

## INTRODUCTION

Stable-isotope labeling with amino acids in cell culture (SILAC)^1,2^ is a powerful tool in quantitative proteomics for comparing biological samples. While chemical labeling strategies like tandem mass tags^3–5^ have increased multiplexing capabilities, SILAC is still widely used for its quantitative accuracy^6^ and its ability to metabolically-encode temporal information^7^. For instance, dynamic SILAC^8,9^ and pulsed-SILAC^10,11^ approaches have become the field standard methods for measuring protein turnover in vivo at a proteome-wide scale.

In a SILAC experiment, proteome mixtures with isotopically-labeled amino acids are analyzed by LC-MS/MS using DDA and the relative quantification of peptide pairs is obtained from the peptide precursor signals in MS1 scans^1^. Comparatively to label-free DDA samples, SILAC samples have better quantitative precision due to decreased technical variation, however SILAC suffers from increased MS1 spectral complexity and redundant sampling of peptides. As a result fewer peptides are quantified^12^.

As an alternative to traditional DDA, data independent acquisition (DIA) strategies^13–15^ with quantification on the MS/MS have also been applied to SILAC samples, improving sampling reproducibility and quantification accuracy. Alternative MS acquisition methods such as BoxCarmax^16^, which leverages a combination of BoxCar^17^, multiplexed MS/MS DIA^18^, and gas phase fractionation^19^, further improve sampling and quantification. However, the wide-window isolation used in most DIA experiments can compromise SILAC quantification, given that asymmetric isolations of the peptide pair can lead to distorted SILAC ratios^20^.

Data-dependent acquisitions that are informed by SILAC peptide pairs have also been developed. For instance, the Mann group demonstrated enhanced quantification of SILAC pairs with poorly defined ratios in the survey MS1 scan by triggering selected ion monitoring scans in real-time^21^. Additionally, the Coon group demonstrated reliable quantification by targeted coisolation and fragmentation of the SILAC peptide pair in direct infusion MS experiments^22^.

Experiments using other stable isotope labeling strategies, such as trypsin-catalyzed ^16^O-to-^18^O-exchange and d0/d3 methyl esterification, have leveraged peptide pairs for identification by comparing heavy and light peptide’s spectra to assist in de novo peptide sequencing^23–25^ and automated peptide search validation^26^. For quantification, Heller et al.^27^ showcased the coisolation of heavy and light ^16^O-^18^O peptide pairs for MS/MS and utilized y-ion fragment pairs for relative quantification. Other methods using chemical^28^ or metabolic^29^ isotopologue labels have demonstrated the feasibility of using isotopically distinct fragment ions for relative MS/MS quantification with accuracy and precision. Analogously, coisolation of SILAC peptide pairs would result in a boost of b-ion fragment MS/MS signal and paired y-ion SILAC MS/MS signal that can be leveraged for database search identification and MS/MS-based quantification.

Here, we implement a MS acquisition method to coisolate SILAC peptide pairs for MS/MS. To analyze coisolated MS/MS, we adapt Comet to perform peptide-spectral matching (PSM) using theoretical spectra of SILAC peptide pairs and we develop a tool to quantify SILAC y-ion pairs from MS/MS spectra. We demonstrate that our method can successfully identify SILAC peptide pairs, while enabling both MS1 and MS/MS-based quantification. We further expand the capabilities of our method to accurately quantify SILAC peptide pairs with overlapping isotopic distributions. Collectively, this work expands our proteomic toolkit for quantitative analysis of SILAC samples.

## METHODS

### Yeast growth and harvest

Two Saccharomyces cerevisiae yeast strains were used to generate SILAC proteome mixtures: DBY10144 and BY4742. Saccharomyces cerevisiae DBY10144 diploid strain (MATa/α) is a lysine prototroph from the FY (S288C) background (parental strains FY3G and FY4H). Saccharomyces cerevisiae BY4742 haploid strain (MATα) is a lysine auxotroph from the FY (S288C) background (parental strain FY2). Two starter cultures (synthetic complete media: 6.7 g/L yeast nitrogen base, 2% glucose, and 2 g/L of drop-out mix with all amino acids except lysine) were grown overnight at 30 °C, one spiked with heavy ^13^C_6_,^15^N_2_-lysine and other with natural isotope abundance (light) lysine (final concentration 0.872 mM for DBY10144 and 0.436 mM for BY4742). Both cultures were diluted using the same media composition (heavy- and light-lysine media respectively) to OD_600_=0.1(DBY10144) and OD_600_=0.05(BY4742). For DBY10144 pellets, yeast cell growth was stalled at OD_600_=~0.75 (~8 doublings with overnight cultures) with 100% trichloroacetic acid (final concentration 10%) and cultures were harvested by centrifugation at 7,000 × g for 10 minutes at 4 °C. Supernatants were decanted and cell pellets were washed with ~10 mL of chilled 100% acetone. Acetone-washed cell pellets were centrifuged at 7,000 × g for 10 minutes at 4 °C, decanted, and cell pellets were snap frozen with liquid nitrogen and stored at −80 °C. For BY4742 pellets, yeast were cultured overnight and harvested at OD_600_=~0.85 (~8 doublings with overnight cultures) by centrifugation at 7,000 × g for 10 minutes at 4 °C. Supernatants were decanted and cell pellets were washed with 2 mL chilled deionized water. Cell pellets were centrifuged at 7,000 × g for 10 minutes at 4 °C, decanted, and cell pellets were snap frozen with liquid nitrogen and stored at −80 °C.

### Cell lysis, protein reduction and alkylation, and protein digestion

Cell pellets were resuspended on ice with 600 μL denaturation buffer composed of 8 M urea, 50 mM Tris pH 8.2, and 75 mM NaCl. Phosphatase inhibitors (10 mM sodium pyrophosphate, 50 mM of sodium fluoride, and 50 mM β-glycerophosphate) were added to the DBY10144 denaturation buffer. Cells were lysed by mechanical agitation using 0.5 mm zirconia/silica beads (60 second bead beating then 90 second rest on ice, repeated four times). Lysates were crudely clarified by centrifugation at 1,200 × g for 1 minute to remove beads followed by cell debris removal via centrifugation at 7,000 × g for 10 minutes at 4 °C. Protein concentration for heavy- and light-lysine lysates was determined by BCA assay (Pierce). Clarified lysates were reduced at 5 mM dithiothreitol (DTT) for 30 minutes at 55 °C, alkylated with 15 mM iodoacetamide in the dark with agitation for 30 minutes at room temperature, and quenched with an additional 5 mM DTT at room temperature for 30 minutes with agitation. Reduced and alkylated samples were diluted 1:1 (v:v) with 50 mM Tris pH 8.9 and 75 mM NaCl to a final pH ~8.5 (DBY10144 samples Tris pH 8.9 buffer contained phosphatase inhibitors: 10 mM sodium pyrophosphate, 50 mM of sodium fluoride, and 50 mM β-glycerophosphate). Lysates were digested with lysyl endopeptidase (LysC; Wako Chemicals in HPLC grade water) 1:100 enzyme to protein substrate ratio overnight at room temperature. LysC digestion was quenched with trifluoroacetic acid (final concentration 1%), and peptides were placed at −80 °C.

### Peptide desalting

Reversed-phase tC18 Sep-Pak columns of 50 mg beads were used to clean up 1.5-1.7 mg of digested yeast lysate. Columns were conditioned with 1 mL methanol and equilibrated with 3 × 1 mL 100% acetonitrile, 1 mL 70 % acetonitrile with 0.25% acetic acid, 1 mL 40% acetonitrile with 0.5% acetic acid, and 3 × 1 mL 0.1% trifluoroacetic acid (TFA). Peptides were loaded twice by gravity. Columns were washed with 3 × 1 mL 0.1% TFA and 1 mL 0.5% acetic acid. Peptides were eluted with 600 μL 40% acetonitrile with 0.5% acetic acid and 400 μL 70% acetonitrile with 0.25% acetic acid. Multiple aliquots of 50 μg or 100 μg peptides were used to generate SILAC mixtures of lysine light to heavy ratios (10:1, 4:1, 2:1, 1:1, 1:2, 1:4, 1:10) for mass spectrometry analysis. All sample aliquots were dried by vacuum centrifugation and stored at −80 °C.

### Liquid chromatography mass spectrometry data acquisition strategies

SILAC peptide mixtures (500 ng) were subjected to nLC-MS/MS on a EASY-nLC 1200 (Thermo Fisher) coupled to a Orbitrap Eclipse Tribrid mass spectrometer (Thermo Fisher). Peptides were loaded on a 100 μm × 3 cm trap column packed with 3 μm C18 beads (Dr. Maisch) and separated using a 90-minute gradient of 80% acetonitrile, 0.1% formic acid on a 100 μm × 30 cm analytical column packed with 1.9 μm C18 beads (Dr. Maisch). Peptides were online analyzed by tandem mass spectrometry using data-dependent acquisition with a cycle time of 3 seconds starting with a full MS1 scan on the Orbitrap over 300-1,500 m/z at 120,000 resolution, normalized automatic gain control set to Standard (100% – 4e5), and injection time set to Auto (max 50 ms). For traditional data-dependent acquisitions, MS/MS was acquired for the most intense m/z precursors (z=2-5) over the 3 second cycle time using the following parameters: dynamic exclusion of 30 seconds, 1.6 m/z isolation window of precursors, HCD fragmentation at 30 normalized collision energy (NCE), 30,000 resolution on the Orbitrap, normalized automatic gain control set to Standard (100% – 5e4), and injection time set to Auto (max 54 ms). For offset left and right wide window scans (Coiso scans) applied to Lys0/Lys8 SILAC mixtures, the same acquisition parameters were used as the DDA MS/MS scan however each triggered precursor was isolated with a 6.5 m/z isolation window, offset −4 Da and 4 Da (left and right respectively). For comparing Coiso scans and DDA scans, the same precursor was subjected to left and right wide window scans and the DDA scan with the same respective parameters as above. For Lys6/Lys8 SILAC mixtures, the isolation offset was set to −1 Da and 1 Da (left and right) with a 5.0 m/z isolation window.

### SILAC peptide pair database search approach and new parameters

The Comet^30,31^ search engine was extended to perform peptide spectral matching on SILAC peptide coisolation theoretical fragments, controlled by the search parameter “silac_pair_fragments”. In addition to specifying whether a standard database search or a coisolation search is performed, this parameter also controls whether to apply the coisolation fragment peaks on both the b- and y-ion series (silac_pair_fragments=1) or only on the y-ion series (silac_pair_fragments=2). Of note, we demonstrated that only paired y-ions should be considered (excluding possible paired b-ions) in peptide-spectral matching (Supp. Discussion) for it standardizes the possible theoretical fragments based on peptide length across all possible candidates, removes a bias towards missed cleaved decoys (Supp. Fig 1a), generates more PSMs (Supp. Fig 1b), and improves E-values (Supp. Fig 1c).

To perform a SILAC coisolation search, a static modification is applied to set the mass of lysine to the light SILAC mass and the mass difference between the light and heavy SILAC reagents is set as a variable modification on lysine residues (e.g. 8.014199 for Lys0/Lys8 or 1.99407 for Lys6/Lys8). All matching candidate peptides are scored by the fast cross-correlation (Xcorr) algorithm^32^.

### Standard E-value and Coisolation E-value for SILAC peptide pair prioritization

Comet calculates an expectation value (E-value) for each reported PSM. For a given PSM, the E-value is defined as the expected number of random peptides that score as well or higher than the PSM’s cross-correlation score (Xcorr). The E-value is calculated based on modeling the incorrect score distribution which, in the case for Comet, is the histogram of random Xcorr scores for each spectrum query searched against candidate peptides from the sequence database. To calculate an E-value, the Xcorr histogram is converted to a cumulative distribution function (CDF) and a linear regression is fit to the right tail of the log10 transform of the CDF. Xcorr scores are extrapolated from the linear regression model to determine each PSM’s E-value^30^.

For a coisolation search, the reported E-value is based on the coisolation Xcorr. Additionally, Comet reports a second E-value (Coiso E-value or e.value_paired) based on Xcorr scores calculated on just the non-triggered fragment ion peaks e.g. only the light fragment ions if the MS/MS spectrum was triggered on a heavy precursor or only the heavy fragment ions if the MS/MS spectrum was triggered on a light precursor. This strategy is similar to calculating the delta Xcorr between the coisolation database search Xcorr (Coiso) and the traditional DDA database search Xcorr (DDA). For each peptide spectral match, all candidate peptides (target or decoy within 20ppm of targeted m/z) will have the delta Xcorr calculated building a delta Xcorr distribution. Similar to calculating a Comet E-value, we can calculate a Coiso E-value based on the expectation value at which the top PSM candidate’s delta Xcorr intersects with the linear regression of the right tail of the −log_10_(ΔXcorr CDF). This score accurately prioritized the correctly coisolated SILAC scan with a drastically higher −log_10_(E-value) than the same precursor incorrectly coisolated. The Coiso E-value score is used to determine when a spectrum has correctly coisolated both light and heavy precursors or incorrectly coisolated only one of the two precursors.

### Database searching S. cerevisiae data with Comet

Prior to database searching, MS raw files were converted to .mzML format using msconvert^33^. The .mzML files from SILAC proteome mixtures were searched with Comet, which can be located at https://sourceforge.net/p/comet-ms/code/1622/tree/branches/release_2019015_silacpair/. The following database search parameters were used: a SGD S.cerevisiae protein sequence database (July 2014, decoys generated for sequence are pseudo-reversed^34^), searching for b-ions, y-ions, and H_2_O/NH_3_ neutral loss fragments, LysC endopeptidase specificity (C-terminal to lysine; max 2 missed cleavages), fixed modification of cysteine carbamidomethyl and [+6.020129] on lysines only when ^13^C_6_-lysine was the light SILAC label, variable modifications of oxidation on methionines, acetylation on protein N-terminus, and heavy lysine delta mass (based on mass difference between light to heavy lysine labels), mass tolerance of 20 ppm for precursor m/z, and mass tolerance of 0.02 Da for fragment ions.The parameter isotope_error was set to 3 for all SILAC ratios used except Lys6/Lys8 SILAC mixtures where isotope_error was set to 1. Coiso SILAC searches were designated silac_fragment_pairs = 1 or 2 (including and excluding paired b-ion fragments respectively, while traditional DDA searches had silac_fragment_pairs=0). Comet generated a pep.xml, pin, and tab delimited text output files for downstream analysis.

### Peptide spectral match (PSM) FDR filtering and MS/MS spectral quantification

We developed a computational Python suite that integrates a variety of publicly available MS software with custom code to generate MS1 and MS/MS quantifications of PSMs from Comet database search results. Using .mzML spectral files, Dinosaur^35^ was used to identify and quantify MS1 spectral features. The Python script takes .mzML files, Comet generated .pin files, and Dinosaur generated .feature.tsv files as inputs. Additionally, Comet generated .pin files could be filtered for PSM results pertaining to correctly coisolated wide window scans (precursors with number of lysines=1), based on Coiso E-value score prioritization, and its corresponding narrow window scan from the same precursor (resultant .pin taken as input). The following steps are conducted using our coisolation Python (v.3.8.1) script: (1) Using Pyteomics^36,37^, relevant spectral header information and MS/MS spectra m/z and intensities are extracted from .mzML files. (2) PSM results from Comet database search engine are FDR filtered at 1% PSM FDR using mokapot^38^. (3) After merging PSM FDR-filtered results with spectral data, we calculate the theoretical masses of b-ion and paired y-ion fragments (z=1 and 2) for the monoisotopic and first and second isotopic peaks. Then, MS/MS peaks were matched to these theoretical masses (maximum intensity observed within 50 ppm tolerance). (4) MS/MS spectral noise was determined using the median of all spectral peaks in the MS/MS scan from the .mzML file^39^. Signal-to-noise ratios for heavy and light fragments were calculated for the average peak intensity of topN, top3, and top6 fragments with quantifiable heavy-light fragment pairs divided by noise. (5) Only annotated fragment isotopes (monoisotopic and/or isotope error peaks with intensity greater than MS/MS spectral noise intensity) that were observed for both heavy and light peptides were considered. Isotope intensities observed in both heavy and light forms were summed to represent the fragment’s heavy and light intensities. (6) MS/MS quantifications were generated as weighted average or median heavy/light ratios using the topN and top2-top6 paired heavy-light lysine paired fragments, excluding y^1+^ fragments. Heavy and light intensities of top3-top6 paired fragment quantifications were fit to a linear regression. A SILAC ratio was extracted based on the slope of the regression with accompanying linear fit coefficient of determination (R^2^). (7) Dinosaur features were mapped to mokapot PSM-filtered results using the following criteria: PSM retention times within the bounds of the peptide MS1 feature, and MS1 feature’s m/z within 50 ppm of the theoretical PSM m/z. If multiple features map to a single PSM, the feature with the max intensity was chosen. (8) All steps were compiled into three output .csv files; one with PSM level summary MS1 and MS/MS quantifications, the second with all annotated heavy and light MS/MS fragments, and the third with annotated pair y-ion fragments only used for quantification. The source code and coiso_silac Python package can be accessed on GitLab at https://gitlab.com/public_villenlab/coiso_silac.

### Lysine6/Lysine8 MS/MS spectral deconvolution and quantification

For ^13^C_6_-lysine (Lys6) and ^13^C_6_,^15^N_2_-lysine (Lys8) SILAC proteome mixtures, the isotopic distributions of coisolated precursor MS1 and MS/MS peptide paired y-ion fragments overlap and require deconvolution for quantification. MS/MS spectral processing and FDR filtering were applied as described above. To determine heavy and light fragment intensity contributions, all paired y-ion fragment intensities were extracted from the MS/MS scan. This resulted in an extended isotopic distribution profile (monoisotopic peak and the first through fourth isotopic peaks) of the light peptide fragment based on the calculated theoretical m/z values. Missing isotope peaks were assigned 0 MS/MS intensity. Light fragment intensities contribute up to all 5 theoretical isotopic peaks, while the heavy fragment intensities only contribute to the second through fourth isotopic peaks. To demarcate the relative contribution of the light and heavy fragments to the second through fourth isotopic peak intensities, the chemical composition of each y-ion fragment for light peptide sequence was determined, and using the Yergey et al. calculation^40^, the theoretical isotopic distribution across the isotopic profile (normalized to the monoisotopic contribution) was calculated.

Using the mathematical approach developed by Chavez et al.^41^, each y-ion fragment’s observed isotopic distribution and the fragment’s theoretical isotopic distribution serve as inputs to compute the fragment’s optimal SILAC ratio (the heavy/light ratio that minimizes the ratio error in their model). Our implementation deviates from Chavez et al.’s approach in that we calculate each y-ion fragment’s theoretical isotope distribution based on each fragment’s molecular composition (compared to using the precursor theoretical distribution for all peptide’s fragments) and we choose a different set of filters for accepting a fragment’s ratio. We applied a filter that the quantification for each fragment can only be calculated if at least two isotopic peaks were observed across the isotopic profile and the heavy and light contributions to the optimal SILAC ratio calculation must be positive values. The PSM’s optimal SILAC ratio and SILAC ratio error were designated by the median of the topN and top3 fragments’ optimal ratio and ratio error for each PSM.

MS1-based quantification was determined via the deconvolution of MS1 precursor signals similarly to MS/MS peptide fragments, using the theoretical distribution of the precursor peptide and observed isotopic distribution for the nearest MS1 or apex MS1 scan for the closest 16 MS1 scans from the triggered MS/MS scan number. Top3 and Top5 MS1 observed isotopic distributions were calculated by summing the isotopic contributions for the 3 or 5 most abundant isotopic distribution signals across subsequent scans. FDR-filtered PSMs with MS1 and MS/MS deconvoluted quantifications were returned as a summary .csv file.

### Data analysis

Spectra for MS1 and MS/MS scans pertaining to figures were directly extracted from .mzML using custom code or Pyteomics^36,37^ and MS/MS spectra were annotated with our custom Coiso annotation code or spectrum_utils^42^. Resultant spectra were plotted in R (version 3.6.1) and RStudio (version 1.4.1103), and all data figures were generated in Adobe Illustrator CS5 (version 15.0.0) and R. All code and data analysis can be accessed via GitLab at https://gitlab.com/public_villenlab/coiso_silac_analysis. All mass spectrometry data and analysis files generated for this manuscript are deposited to the ProteomeXchange Consortium by the PRIDE partner. The PRIDE project identification number is PXD033016, and the reviewer username is; password oQ7OiEl7.

## RESULTS

### Rationale for coisolation SILAC acquisition and database searching

To coisolate SILAC pairs, we use a DDA approach where the detected precursor m/z’s are isolated in a 6.5 m/z-wide isolation window that is offset to the center of the SILAC pair. For a light precursor the offset would be set to the right and for heavy precursors to the left. Our main goal here is to provide proof of principle of the coisolation approach. To avoid relying on the instrument designating precursors as light or heavy, we acquire MS/MS for both offset isolations, and compare to standard DDA MS/MS (i.e. with 1.6 m/z-wide isolation window) for the same precursor (Fig 1a). We adapted Comet to be able to assign peptides from MS/MS spectra of coisolated SILAC pairs (Fig 1b). In the manuscript, we refer to the combination of the wide window MS/MS acquisition and the coisolation Comet search as “Wide Coiso”, and the standard DDA MS/MS and Comet search as “Narrow DDA”.

**Figure 1:**
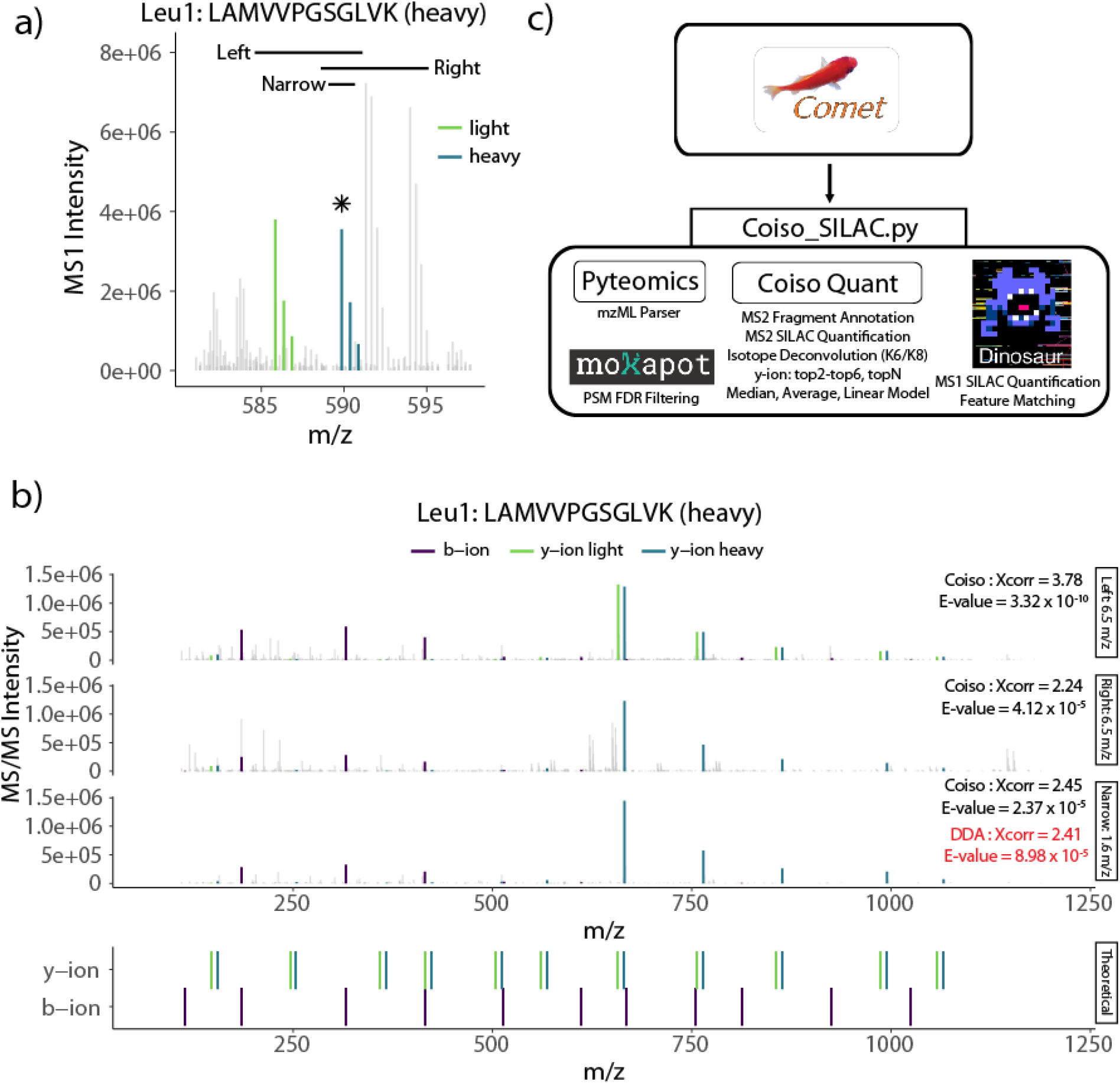
Coiso SILAC computation workflow and MS acquisitions. **a)** From the MS1 scan, Coiso 6.5-m/z wide isolation window (offset −4 Da left and +4 Da right) and DDA narrow window 1.6 m/z isolation window for a triggered heavy SILAC peptide from Leu1 (asterisk). Coiso MS acquisition aims to capture SILAC peptide pairs (light (green) and heavy(blue)) for fragmentation and MS/MS. **b)** MS/MS of left and right Coiso scans and narrow window DDA scan for the triggered Leu1 peptide in (**a**). Comet’s E-values and Xcorr for the Coiso search are in black and standard DDA search are in red. Theoretical fragments for Coiso search are in the bottom panel at theoretical m/z values. Peptide spectral matched b-ions (purple), light y-ions (green), and heavy y-ions (blue) are highlighted and other MS/MS peaks are grey-scaled. **c)** Comet database searching for Coiso SILAC MS acquisitions performs peptide spectral matching to SILAC peptide paired fragments designated by the parameter “silac_pair_fragments”. PSMs are matched to Pyteomics-parsed MS/MS spectral data, FDR-filtered with mokapot, mapped to Dinosaur MS1 features, and PSM y-ion paired fragments are annotated and quantified.

We expect the coisolation and analysis of peptide pairs (Wide Coiso) will yield three improvements. First, fragmentation spectra of coisolated SILAC peptide pairs feature merged b-ions and split y-ions (Fig 1b top panel). In a Comet search, the increased ion representation should increase Xcorr values (Fig 1b). Second, to overcome the increased complexity in a wide isolation window, we apply a narrow mass tolerance to increase the specificity of the precursor. We expect these two features will prioritize the true pair over other candidates and paired decoys, thus improving Comet E-values and overall identifications. Third, quantification can be performed in the MS/MS using the y-ions, which are expected to have higher signal-to-noise ratios than precursor signals^43^.

Here, we assessed the performance of the method by comparing peptide-spectral matching metrics and identifications between SILAC pair MS/MS and DDA MS/MS for the same precursors. Also, we evaluate quantification precision and accuracy for MS1 precursor and MS/MS y-ion pair features.

### Comet database search and quantification tool for SILAC peptide pairs

To analyze SILAC pair data, a two step analysis pipeline was generated. First, Comet was adapted to perform peptide spectral matching (PSM) for SILAC pairs using heavy and light fragments. This feature is enabled by the “silac_pair_fragments” parameter. For all candidate targets and decoys within the precursor mass tolerance, theoretical paired spectra are generated and matched to calculate cross correlation (Xcorr) scores^32^. With calculated Xcorr values, PSMs can be assigned an E-value, or an expectation score, as a metric to standardize the confidence of the reported PSMs and calibrate Xcorr values across scans. E-values serve as an effective single metric to differentiate target and decoy PSMs^30^, thus enabling FDR-based PSM filtering to establish a high confidence set of identifications for downstream analysis.

The second component of the analysis pipeline is an open source Python package, coiso_silac, that performs SILAC quantification with coisolation data. In the package, we integrate the publicly available software mokapot^38^ to FDR filter Comet PSMs, Pyteomics^36,37^ to extract MS spectra m/z and intensity, and Dinosaur^35^ to identify MS1 precursor features. With this information, we annotate peptide fragments in MS/MS spectra, calculate MS/MS-based SILAC ratios, and map MS1 features to PSMs (Fig 1c). MS/MS SILAC ratios are generated for a variety of different summation strategies and variable number of most abundant topN paired y-ions. This quantification pipeline generates a result file containing 1% FDR filtered PSMs with scoring metrics and MS1 and MS/MS quantifications for downstream analysis.

### Coisolating SILAC pairs improves identification metrics compared to DDA

We applied the MS isolation schema from Fig. 1a to analyze seven S. cerevisiae proteome samples mixed at different SILAC ratios (10:1, 4:1, 2:1, 1:1, 1:2, 1:4, and 1:10; light:heavy; Lys0:Lys8). For simplicity, initial coisolation MS acquisitions were designed based on SILAC peptides with a single lysine. Thus, peptide identifications that did not have exactly one lysine were removed from the analysis. Since both wide offset MS/MS scans are analyzed for each precursor, only the wide offset scan containing the SILAC pair was compared to the matching DDA MS/MS scan (See Methods).

We assess method feasibility by comparing identification metrics between our Wide Coiso approach and traditional Narrow DDA for matching precursors. Herein, the metrics for PSM spectral representation (Xcorr) and confidence in PSM assignment (E-value) were compared. In the 1:1 SILAC labeled sample, the Wide Coiso approach showed higher Xcorr scores for 96% of the PSMs compared to Narrow DDA (Fig 2a), likely due to the additional, paired y-ions and intensity increase in b-ion signals. To benefit from this increase in Xcorr with Wide Coiso in the PSM filtering step, we should observe a greater increase for true matches than for false matches. Indeed, we observed the standardized E-value metric showed a 42% improvement for Wide Coiso compared to Narrow DDA for matching PSMs at 1% FDR (Fig 2b).

**Figure 2:**
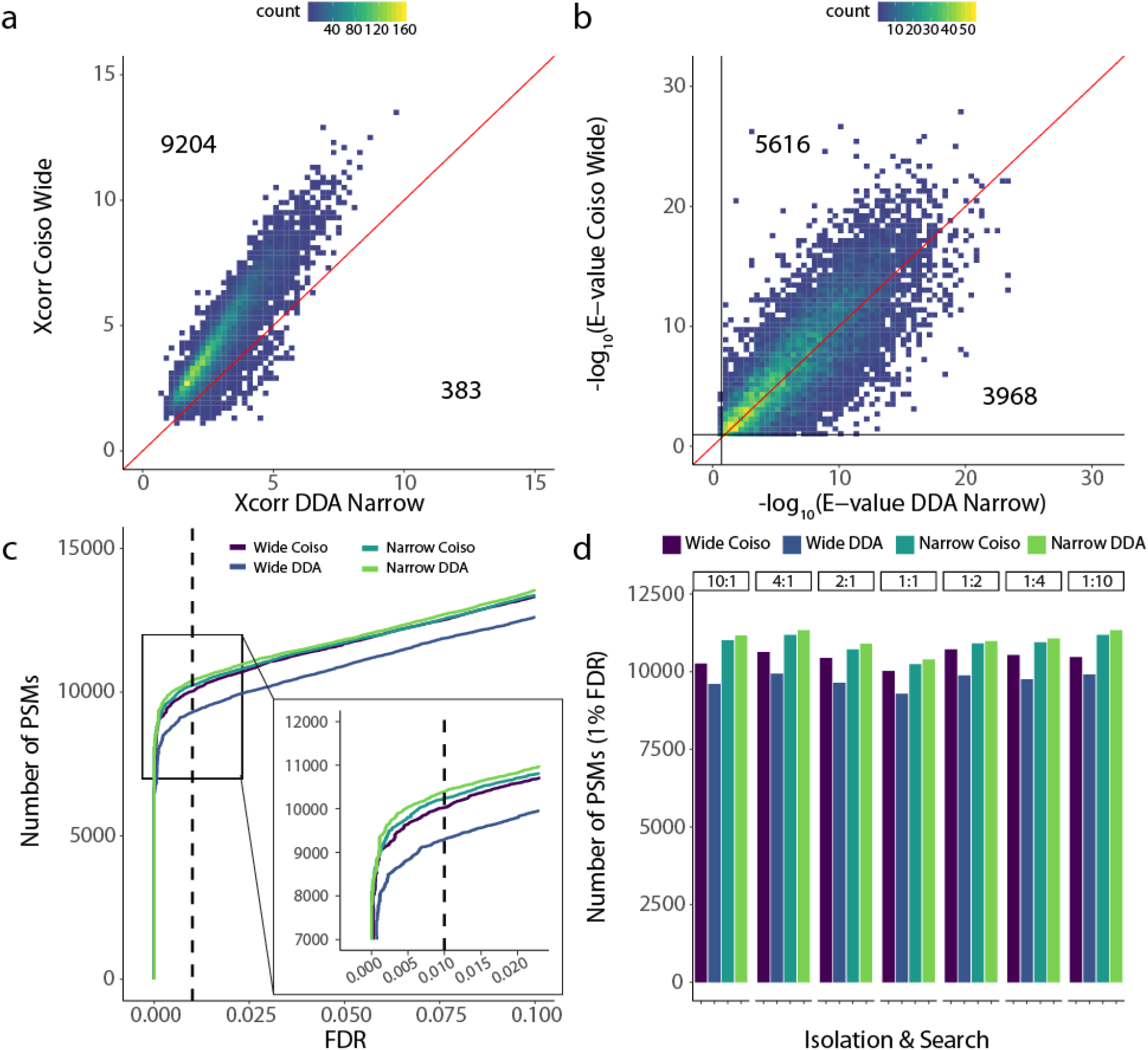
Coiso SILAC Comet identification performance is comparable to traditional DDA. **a)** Density-binned scatter plot comparison of Xcorr from correctly coisolated wide window scan searched with Coiso algorithm and narrow window scan searched with DDA parameters for matching targeted precursor for 1:1 *S. cerevisiae* SILAC mixture. **b)** Same as (**a**) for Comet E-values. **c)** Number of PSMs at specified FDR using E-value target-decoy competition for all MS acquisition and search combinations. The combinations consider correctly coisolated Coiso scans and narrow window scans searched with Coiso or traditional DDA Comet search parameters for 1:1 *S. cerevisiae* SILAC mixture. Inlet zooms over an FDR = 0.01 cut-off. **d)** PSM identifications at 1% FDR based on Comet E-values for PSMs with one lysine across the scan acquisition:Comet search parameter combinations in **(c)** for seven *S. cerevisiae* SILAC ratio proteome mixtures.

Across the run, Wide Coiso and Narrow DDA showed similar E-value distributions for both decoy hits and target hits (Supp. Fig 2a). However, we observed a shifted distribution toward higher −log_10_(E-values) in the Wide Coiso approach (Supp. Fig 2b,c).

Comet E-values are often used to filter PSMs to a defined FDR and generate a high-confidence set of PSMs for downstream analysis. We assessed the filtering capabilities of the E-value metric for Coiso Wide and Narrow DDA by estimating PSM false discovery rate (FDR) using target-decoy competition. In the 1:1 SILAC sample, we observed that the target/decoy discriminatory capability of Wide Coiso E-values is similar to that of Narrow DDA E-values, however Narrow DDA is slightly more sensitive (Fig 2c). When comparing total PSM identifications at 1% FDR across SILAC mixtures, Wide Coiso captures 92% for 1:10 and 10:1, 94% for 1:4 and 4:1, 97% for 1:2 and 2:1, and 97% for the 1:1 SILAC ratio samples compared to Narrow DDA (Fig 2d). Thus, we conclude that our coisolation pair method provides a similar capability to identify SILAC peptide pairs proteome-wide as DDA.

Additionally, we searched the Wide and Narrow MS acquisitions with the opposing Comet search DDA and Coiso parameters, respectively. Not surprisingly, Narrow Coiso slightly underperformed Narrow DDA because the data does not contain paired fragments. Wide DDA had the lowest number of identifications because the data has paired fragments that increase spectral complexity without benefit to the analysis.

### Coisolation score for identifying scans that coisolate SILAC pairs

While our coisolation approach generally improves PSM E-values, E-values alone are not strong predictors of whether the SILAC peptide pair is coisolated. Indeed, we observed that wide window isolations that exclude one of the SILAC features can generate a PSM E-value that passes the PSM-level 1% FDR filter (Fig 1b 2nd panel: right isolation). Therefore, we sought to generate a coisolation score that could prioritize wide window MS/MS that successfully coisolate the SILAC pair.

If a SILAC pair is coisolated, we would expect to see y-type fragments from both the targeted and the non-targeted SILAC precursor. To generate a coisolation score, we first calculate a Xcorr considering only the y-ions of the non-targeted SILAC precursor, which are only present during coisolation. Then, we compare this Xcorr against all candidates within the MS1 ppm tolerance to calculate an expectation score or Coiso E-value (Supp. Fig 3). Our Coiso E-value should calibrate the non-targeted precursor’s y-ion Xcorr contribution across scans and standardize our confidence in SILAC pair detection.

To test whether the Coiso E-value can correctly prioritize coisolated SILAC peptide pairs, we analyzed a S. cerevisiae proteome mixture with a 1:1 SILAC sample using only the wide isolation MS acquisitions from Fig 1a. Since both wide offset MS/MS scans are analyzed for each precursor, −log_10_(Coiso E-values) should be higher for coisolated wide scans compared to the respective non-coisolated scan. Coisolated wide scans are defined as targeting heavy precursors with left offset and light precursors with right offset. With the 6.5 m/z wide windows, coisolation can occur for all charge states when one lysine is present and charges z=4-6 when two lysines are present. All other lysine-charge combinations will not coisolate for both offset MS/MS, as the non-targeted precursor will fall outside the MS1 isolation window.

We found that coisolated scans yield better standard Comet E-values, but this increase in score performance alone is not enough to distinguish these scans from non-coisolated scans (Supp. Fig 4).

In contrast, our new Coiso E-value successfully classified coisolated and non-coisolated SILAC pair spectra (Fig 3). As expected, this separation is dependent on the number of lysines and charge of the peptide precursor, which determine the SILAC pair m/z separation and whether a coisolation is possible with the 6.5 m/z isolation window used in this study. Thus, precursors with no coisolation have Coiso E-values similar to those of decoy hits (2 Lys with z=2-3, and 3 Lys with z=2-6). Precursors with coisolation in only one offset MS/MS separate between non-coisolated and coisolated scans with the expected heavy or light peptide assignments (z=2-3 & 1 Lys; z=3-6 & 2 Lys). Lastly, precursors that coisolate with both offset MS/MS demonstrate Coiso E-values along the diagonal (z=4-6 & 1 Lys) (Fig 3). Thus, our Coiso E-value can serve as a useful predictor for correctly coisolated SILAC peptide pairs.

**Figure 3:**
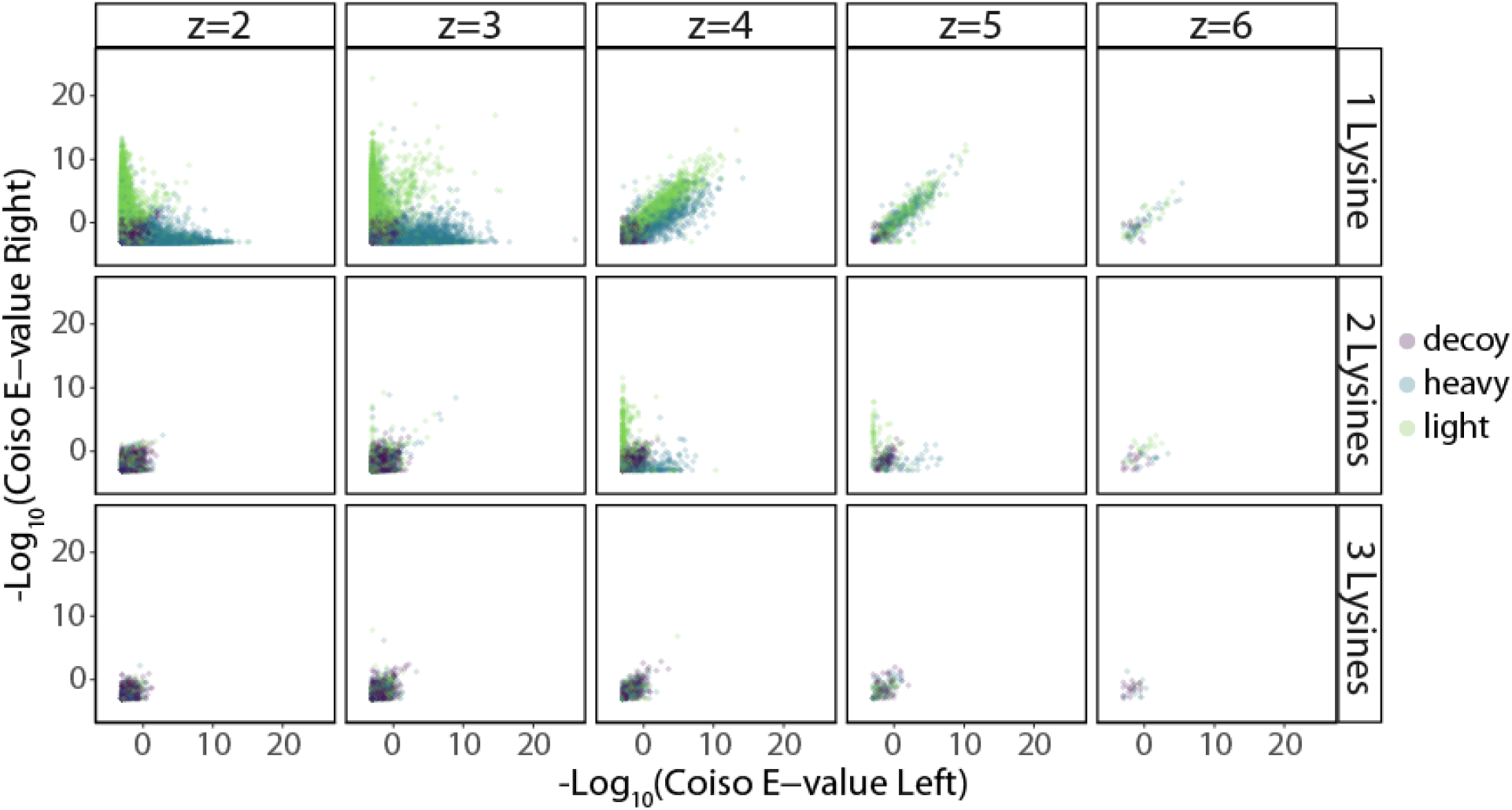
Coiso E-value for prioritizing SILAC peptide pair coisolation and predictive quantifiability. Scatterplot for Coiso E-value of the 1:1 *S. cerevisiae* SILAC proteome mixture faceted by PSM charge state and number of lysines for matching left and right offset Coiso scans. PSM assignment either heavy (blue), light (green), or decoy (purple) based on correctly Coiso scans PSM sequence.

Because Coiso E-values are a metric for robust paired y-ion signals, it can be further applied to filter PSMs for high-quality quantifications, providing FDR-control of SILAC quantifications.

### MS/MS quantification of SILAC peptide pairs

To assess quantification performance, we analyzed seven SILAC S. cerevisiae proteome mixtures (labeled with Lys0 and Lys8) using Wide MS/MS with both offsets. PSMs were filtered for the offset with the higher −log_10_(Coiso E-value) and peptides with one lysine (See Methods). For each PSM, we calculated SILAC ratios at the MS1 level using precursor signals and at the MS/MS level using the most abundant y-ion signals. The number of quantified PSMs at the MS/MS level and its overlap with quantified MS1 precursors vary based on the the number of top y-ions used for quantification (Supp. Fig 5-6). Here, we highlight MS/MS quantification using the Top3 and Top4 most abundant y-ions, since they showed the best balance between the number of PSMs quantified and quantification accuracy and precision.

Due to the increased MS/MS sensitivity achieved by precursor isolation, we expected more quantifications at the MS/MS level compared to the MS1 level. In a 1:1 Lys0:Lys8 S. cerevisiae proteome mixture, we observed a 14% increase in quantified PSMs by Top3 y-ion pair MS/MS compared to MS1 (Fig 4a). This increase was further amplified as the SILAC ratio deviated from 1:1. We observed a 15% improvement with the 1:2 and 2:1 mixtures, 23% improvement with 1:4 and 4:1, and 40-51% improvement with 1:10 and 10:1 (Fig 4a). MS1 and MS/MS quantifications using the Top3 and Top4 y-ion pairs largely overlapped (Fig 4b-c).

**Figure 4:**
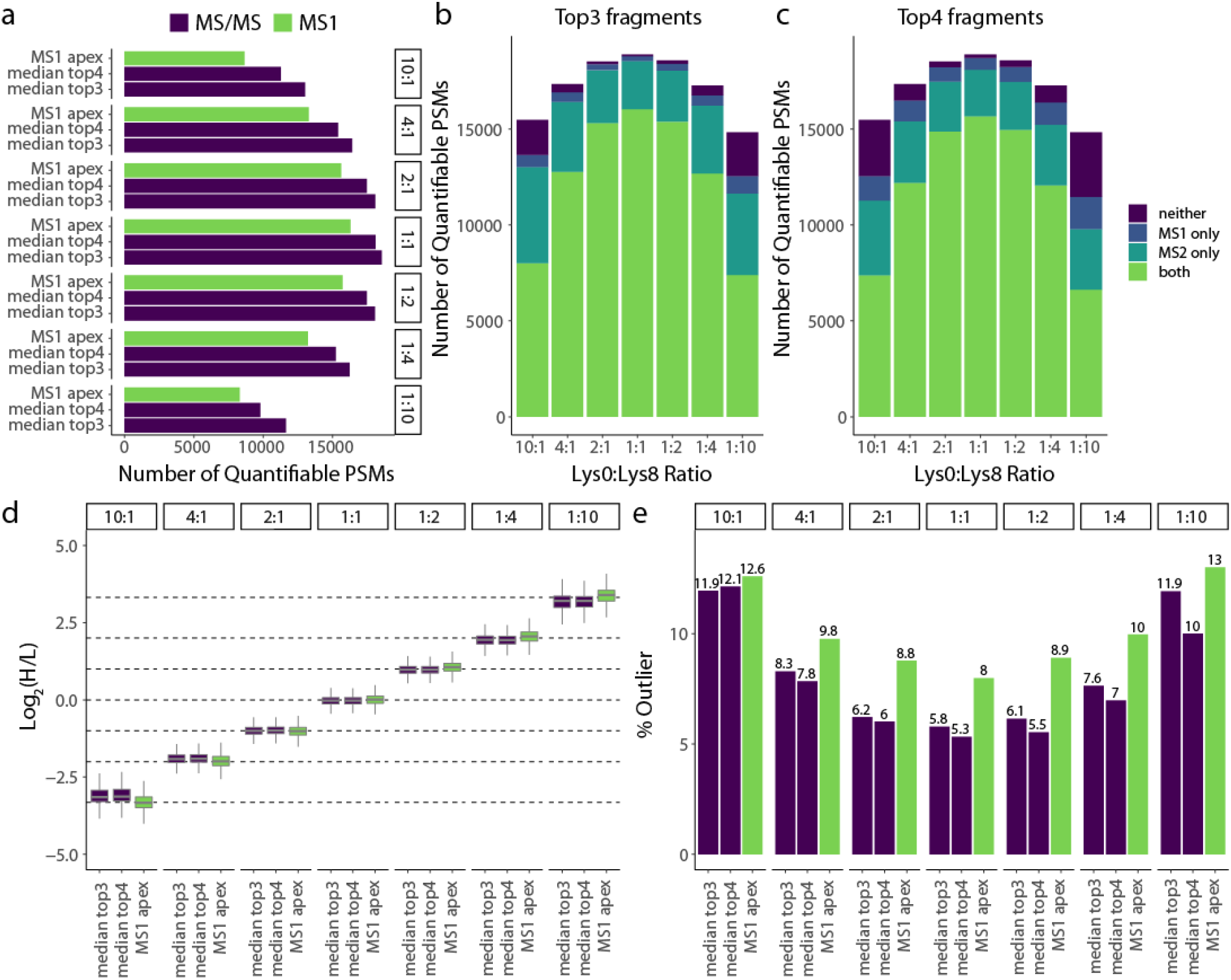
Coiso SILAC enables more quantifications with improved precision and less outliers. **a)** Barplot of the number of quantifiable PSMs for MS1 (green) and the two best Coiso MS/MS quantification methods (purple). Plots consider the same set of PSMs for each of the SILAC *S.cerevisiae* proteome mixtures (Lys0:Lys8 ratio respectively). **b-c)** Number of quantified PSMs across the same samples as in (**a**) depicted as a stacked barplot outlining the overlap between MS1 quantifications and MS/MS quantifications considering top3 (**b**) or top4 (**c**) quantifiable y-ion fragment MS/MS-based quantification filter. Overlap designations are the following: both: quantifiable MS1 and MS/MS (green); neither: not quantifiable in MS1 and MS/MS (purple); MS1 only (dark blue); MS/MS only (light blue). **d)** Box plots for peptide-spectral matches for the same samples as in (**a**). MS/MS-based quantification methods (purple) filtered for top3 or top4 quantifiable paired y-ion fragments. MS1-matched SILAC features from Dinosaur (green) using either apex or sum-based quantification. Box plots represent the distribution from all quantifiable PSMs from (**a**). In the box plot, the horizontal line represents the median, box designates the IQR, and the whiskers indicate 1.5 × IQR from the box ends. **e)** Bar plots of percentage peptide-spectral matches that are outliers from the SILAC *S.cerevisiae* proteome mixture distributions (Lys0:Lys8 respectively) as in (**a**). MS/MS quantification methods (purple) filtered by top3 or top4 quantifiable paired y-ion fragments and Dinosaur MS1-derived quantifications (green) are represented here. Outliers are PSM log_2_(ratios) > Q3 + (1.5 × IQR) or log_2_(ratios) < Q1 – (1.5 × IQR).

We explored multiple ways to calculate SILAC ratios at the MS/MS level using the sum, median, or linear regression across a number of TopX y-ion pairs (Supp Fig 7). For Top3 and Top4 y-ion pairs, median-based MS/MS quantifications showed narrower distributions than MS1 quantifications (Fig 4d), suggesting improved precision. For 1:1, 1:2, and 2:1 Lys0:Lys8 SILAC mixtures, the distributions of heavy-to-light ratios for MS/MS quantifications accurately center at the expected values. However, SILAC ratios further from 1:1 showed mild (1:4 and 4:1) to moderate (1:10 and 10:1) ratio compression. The extent of ratio compression did not correlate with MS/MS fragment ion signal, and the relationship between SILAC ratios and MS intensities were similar between MS1 and MS/MS quantifications (Supp. Fig 8).

Due to MS1 complexity, many precursor signals fall close to the noise level, resulting in poor quantifications. We expect that our coisolation method’s gas phase enrichment would improve signal-to-noise resulting in fewer outlier quantifications. To compare, we define an outlier as a PSM quantification that lies 1.5 times the interquartile range (IQR) away from the first and third quartile. The percentage of outlier PSMs varies by MS/MS quantification approach and top y-ion pair filters (Supp. Fig 9). Remarkably, for all SILAC ratios, Top3 and Top4 median-based MS/MS quantifications had fewer outliers compared to MS1 quantifications (Fig 4e).

### Quantifying SILAC peptide pairs with overlapping isotopic distributions

An avenue to improve the SILAC coisolation method is to increase its multiplexing capacity. This can be achieved by combining isotopically labeled amino acids with smaller delta masses, while maintaining a similar size isolation window. MS1-based quantification of SILAC mixtures with overlapping isotopic distributions are challenging to deconvolute using traditional DDA. The coisolation method could improve SILAC quantification due to reduced isotopic overlap of peptide fragment ions in the MS/MS. Thus, we set out to test the coisolation method for proteomes with SILAC labels separated by 2 Da, potentially enabling a 5-plex over a 8 Da range.

We generated ^13^C_6_-lysine (Lys6) and ^13^C_6_,^15^N_2_-lysine (Lys8) labeled S. cerevisiae proteome mixtures spanning seven ratios (10:1, 4:1, 2:1, 1:1, 1:2, 1:4, and 1:10 Lys6:Lys8). We applied our coisolation method using 1 Da left and right offsets with 5 m/z wide isolations. PSMs were assigned using the version of Comet we adapted for SILAC pairs and filtered for the offset MS/MS with a higher −log_10_(Coiso E-value) and one lysine (See Methods).

In Lys6:Lys8 SILAC mixtures, the second isotope of the lighter (Lys6) precursor of the SILAC pair has the exact same mass as the monoisotopic peak of the heavier (Lys8) precursor, aggregating their signals. This overlap continues for heavier isotopes of the distribution. We deconvoluted signals for precursors and y-type fragments using the mathematical approach in Chavez et al.^41^. This approach fits the theoretical isotopic distribution^40^ to the observed isotopic spectra to calculate an optimal SILAC ratio (R_opt_) (Fig 5a).

**Figure 5:**
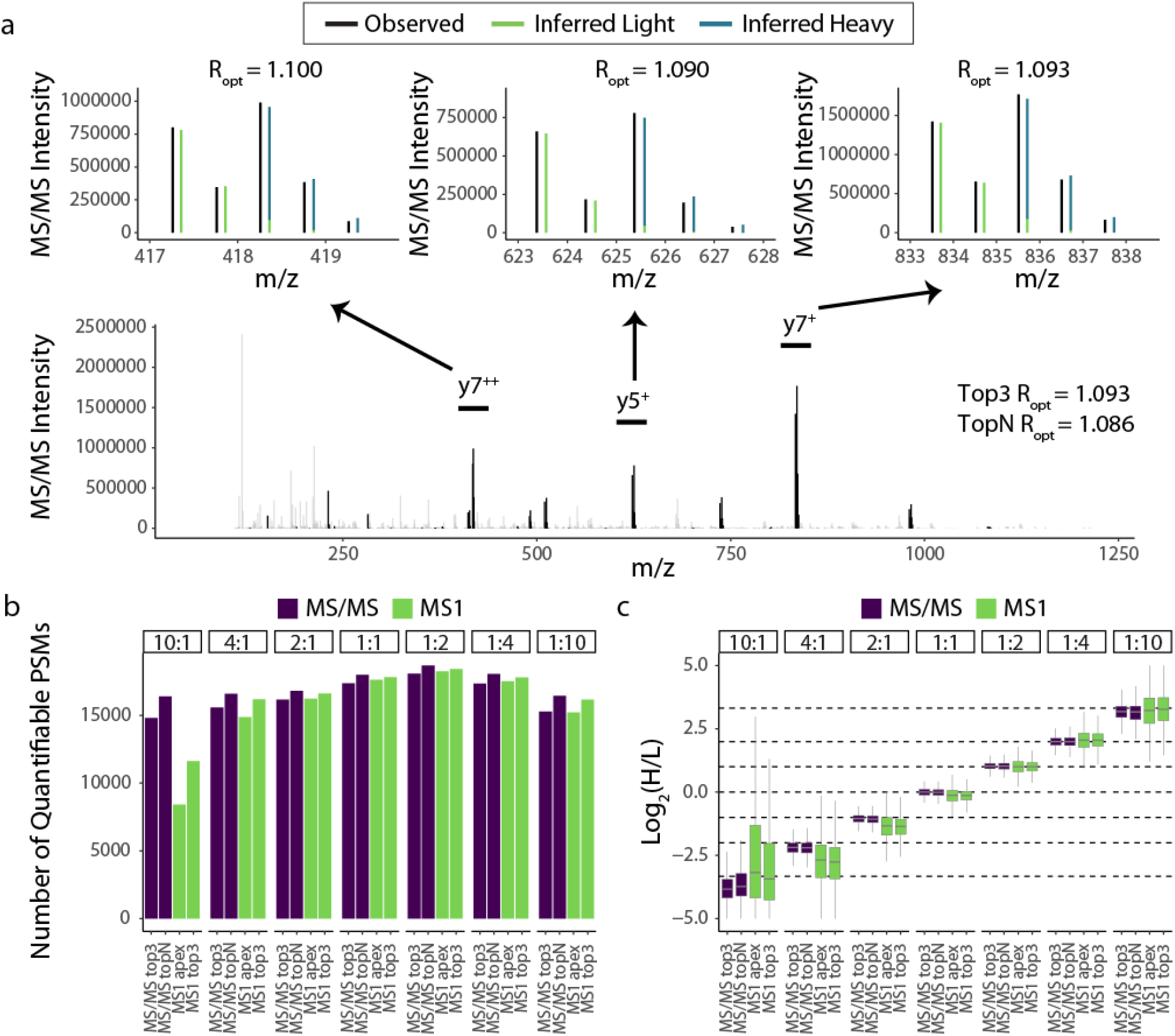
Coiso SILAC of Lys6:Lys8 mixtures via isotope deconvolution for robust MS/MS quantifications. **a)** Coiso SILAC MS/MS spectra (bottom panel) of PSA1 ETFPILVEEK light peptide with peptide fragments (black) and unannotated peaks (grey). Top3 peptide paired fragments (y7^++^: left, y5^+^: middle, y7^+^:right) zoomed in over the fragment’s observed isotope distribution (black) and the inferred light (green) and inferred heavy (blue) fragment intensity contributions based on the model-defined optimal SILAC ratio (R_opt_ = Log_2_(heavy/light)). **b)** Bar plot of number of quantifiable PSMs for topN and top3 fragment ion filter requirement (faceted by proteome mixtures defined by SILAC Lys6:Lys8 ratios). MS1 quantifications based on precursor pair signals in apex MS1 scan or three most intense successive MS1 scans. **c)** Box plots for the distribution of all quantifiable PSMs from (**b**). MS/MS-based quantification methods (purple) filtered for top3 or topN quantifiable paired y-ion fragments. In the box plot, the horizontal line represents the median, box designates the IQR, and the whiskers indicate 1.5 × IQR from the box ends.

For MS/MS-based quantification, we applied this deconvolution approach to calculate R_opt_ ratios for each y-ion fragment and then calculated a SILAC ratio for each PSM as the median R_opt_ among the top3 and topN most abundant y-ion pairs. For MS1-based quantification, we applied the deconvolution method for overlapping precursor signals, generating a R_opt_ for the chromatographic peak apex and a median R_opt_ for the three most intense successive MS1 scans (See Methods).

The deconvolution approach resulted in similar numbers of quantified MS1 and MS/MS features in Lys6:Lys8 samples (Fig 5b). In mixtures 4:1 through 1:4, we observed a similar number of quantified PSMs based on y-ion pairs for Lys6:Lys8 and Lys0:Lys8 (Fig 5b, Supp. Fig 10a). However, at the most extreme ratios, more PSMs were quantified by MS/MS in Lys6:Lys8 samples compared to Lys0:Lys8 samples (Supp. Fig 10a). Surprisingly, the Lys6:Lys8 mixture with the most quantified PSMs was the 1:2 Lys6:Lys8 ratio, possibly due to the asymmetrical light isotopic contribution boosting the heavy precursor signal.

We observed Lys6:Lys8 MS/MS-based quantifications accurately mapped to the expected SILAC ratios (Fig. 5c) and had similar precision to Lys0:Lys8 mixtures (Supp. Fig 10b). In 1:4 and 1:10 Lys6:Lys8 SILAC mixtures, we observe mild SILAC ratio compression, while the 4:1 and 10:1 SILAC mixtures deviate towards more extreme ratios. This is likely due to the limitations to the isotopic deconvolution, which is supported by our observation of ratios being more extreme for shorter peptides (i.e. less overlap between isotopic envelopes) in the 10:1 Lys6:Lys8 sample (Supp. Fig. 11).

Collectively, we demonstrate the coisolation method can reliably and precisely quantify proteome mixtures containing SILAC labels with overlapping isotopic distributions.

## DISCUSSION

Here we developed a SILAC coisolation analysis platform equipped with novel MS/MS acquisition schema, database searching strategy, and quantification pipeline. The offset, wide window MS isolations enable coisolation of SILAC peptide pairs for joint MS/MS analysis. The adapted Comet search for SILAC pairs leverages light and heavy fragments for peptide-spectral matching. The coisolation method improves Comet identification scores and achieves a similar number of PSMs compared to traditional DDA. We generated a Coiso E-value score that prioritizes MS/MS with successfully coisolated peptide pairs and enables FDR control for SILAC quantification. Furthermore, our coisolation method offers MS/MS-based quantification with little to no interference, using y-ion pairs. Quantification of coisolated SILAC pairs outperformed MS1 quantification in the number of quantified PSMs, quantification precision, and percentage of outlier quantifications. However, the method suffered from some ratio compression and reduced accuracy at extreme SILAC ratios. Of note, the coisolation method provides both, standard MS1 quantifications and the additional MS/MS quantifications.

Another advantage of the coisolation method is the capability to deal with labeling schemes that produce overlapping isotopic envelopes. Here we show the performance of our method with Lys6:Lys8 labeling and the method can be applied to labeling schemes that combine Lys0:Lys2:Lys4:Lys6:Lys8, increasing sample multiplexing.

The two main limitations of this study are related to the current coisolation MS/MS acquisition implementation. First, we performed wide window MS/MS for both offsets to ensure coisolation in at least one of the offsets. This duplicates the number of scans per SILAC precursor pair and slows down acquisition. Second, much like in data independent acquisition, wide window isolations result in high complexity MS/MS, which can be difficult to match to a peptide sequence. These two limitations result in fewer peptide identifications than using traditional DDA.

The main goal of our study was to demonstrate the feasibility of the coisolation method when the SILAC peptide pair is successfully coisolated. In future work, we will explore strategies to decrease spectral complexity and improve proteome sampling in combination with the coisolation method. For example, we can apply offline HPLC fractionation to reduce sample complexity. Also, online ion mobility separations (IMS) could separate SILAC peptide pairs by their collisional-cross section (CCS). DIA-SIFT^44^, which couples SILAC-DIA with IMS, demonstrated robust precision and accuracy for SILAC MS/MS quantification, suggesting possible advantages when applying IMS to the coisolation method.

With programmatic accessibility to the mass spectrometer^45^, we envision greater improvements to the coisolation method. We propose implementing an acquisition workflow in which MS/MS scans are triggered on SILAC pair MS1 features detected in real-time, with dynamic adjustment of the center and width of the isolation window. This adjustment will accelerate acquisition by eliminating the need to acquire both offset MS/MS scans for each precursor. Additionally, to increase sampling and reach proteome depths similar to label-free approaches, dynamic exclusion could be programmed at the level of the SILAC pair rather than individual precursors. Finally, sequential isolation of paired light and heavy precursors with an MSX approach^18^ can minimize the MS/MS spectral complexity while increasing SILAC pair signals. Collectively, these developments can substantially improve the coisolation method maximizing the number of quantified peptides achieved per run.

In future, we envision the coisolation SILAC method to offer unique advantages to MS analysis of phosphoproteomes. During peptide-spectral matching, site diagnostic fragment ions that distinguish between phosphate localizations must be observed in order to precisely localize the phosphosite. We expect that the paired y-ions collected in the MS/MS of the coisolation method will offer additional site diagnostic fragments and increase the confidence in site localization assignments. Additionally, site diagnostic ions could be used in the coisolation method to separately quantify phosphopeptide positional isomers, which is not possible if the quantification is done at the MS1 level.

This study presents a coisolation MS method that can successfully identify and quantify SILAC labeled proteomes by isolating and cofragmenting peptide pairs. This approach expands our proteomics toolkit for analyzing SILAC samples and offers exciting new opportunities for future development.

## AUTHOR CONTRIBUTIONS

I.R.S., J.K.E., R.A.R.-M., and J.V. conceived the study and designed the experiments. I.R.S. conducted the experiments with assistance from A.H. and advice from J.K.E., R.A.R.-M., A.S.B., and J.V. J.K.E. developed the Comet database search for paired MS/MS with advice from I.R.S. I.R.S. developed the Python quantification package (coiso_silac) with advice from A.S.B. and J.K.E. A.L. helped with streamlining method development. I.R.S. analyzed the data with advice from A.S.B. and J.K.E. J.V. supervised the study. I.R.S., J.K.E., and J.V wrote the paper, and all authors edited it.

## ACKNOWLEDGEMENTS

We would like to thank the members of the Villén Lab for scientific discussions pertaining to the project, particularly Bianca Ruiz, Kyle Hess, and Mario Leutert. Also, we would like to thank Will Fondrie, Andy Keller, Bill Noble, and Devin Schweppe for valuable discussions pertaining to the method’s development. I.R.S. was supported by the NIH training grant T32HG000035. A.S.B. was supported by the NIH training grant T32LM012419. Most of this work was supported by the NIH grant R35GM119536 to J.V. The Villén lab is additionally supported by NIH grants R01AG056359, R01NS098329, and RM1HG010461, Human Frontiers Science Program grant RGP0034/2018, and a Research Program grant from the W.M. Keck Foundation. This work is also supported in part by the University of Washington Proteome Resource UWPR95794.

## REFERENCES

(1) Ong, S.-E.; Blagoev, B.; Kratchmarova, I.; Kristensen, D. B.; Steen, H.; Pandey, A.; Mann, M. Stable Isotope Labeling by Amino Acids in Cell Culture, SILAC, as a Simple and Accurate Approach to Expression Proteomics*. Mol. Cell. Proteomics 2002, 1 (5), 376–386. https://doi.org/10.1074/mcp.M200025-MCP200.

(2) Blagoev, B.; Kratchmarova, I.; Ong, S.-E.; Nielsen, M.; Foster, L. J.; Mann, M. A Proteomics Strategy to Elucidate Functional Protein-Protein Interactions Applied to EGF Signaling. Nat. Biotechnol. 2003, 21 (3), 315–318. https://doi.org/10.1038/nbt790.

(3) Thompson, A.; Schäfer, J.; Kuhn, K.; Kienle, S.; Schwarz, J.; Schmidt, G.; Neumann, T.; Hamon, C. Tandem Mass Tags: A Novel Quantification Strategy for Comparative Analysis of Complex Protein Mixtures by MS/MS. Anal. Chem. 2003, 75 (18), 4942–4942. https://doi.org/10.1021/ac030267r.

(4) McAlister, G. C.; Huttlin, E. L.; Haas, W.; Ting, L.; Jedrychowski, M. P.; Rogers, J. C.; Kuhn, K.; Pike, I.; Grothe, R. A.; Blethrow, J. D.; Gygi, S. P. Increasing the Multiplexing Capacity of TMTs Using Reporter Ion Isotopologues with Isobaric Masses. Anal. Chem. 2012, 84 (17), 7469–7478. https://doi.org/10.1021/ac301572t.

(5) Li, J.; Van Vranken, J. G.; Pontano Vaites, L.; Schweppe, D. K.; Huttlin, E. L.; Etienne, C.; Nandhikonda, P.; Viner, R.; Robitaille, A. M.; Thompson, A. H.; Kuhn, K.; Pike, I.; Bomgarden, R. D.; Rogers, J. C.; Gygi, S. P.; Paulo, J. A. TMTpro Reagents: A Set of Isobaric Labeling Mass Tags Enables Simultaneous Proteome-Wide Measurements across 16 Samples. Nat. Methods 2020, 17 (4), 399–404. https://doi.org/10.1038/s41592-020-0781-4.

(6) Hanke, S.; Besir, H.; Oesterhelt, D.; Mann, M. Absolute SILAC for Accurate Quantitation of Proteins in Complex Mixtures Down to the Attomole Level. J. Proteome Res. 2008, 7 (3), 1118–1130. https://doi.org/10.1021/pr7007175.

(7) Mann, M. Functional and Quantitative Proteomics Using SILAC. Nat. Rev. Mol. Cell Biol. 2006, 7 (12), 952–958. https://doi.org/10.1038/nrm2067.

(8) Pratt, J. M.; Petty, J.; Riba-Garcia, I.; Robertson, D. H. L.; Gaskell, S. J.; Oliver, S. G.; Beynon, R. J. Dynamics of Protein Turnover, a Missing Dimension in Proteomics. Mol. Cell. Proteomics 2002, 1 (8), 579–591. https://doi.org/10.1074/mcp.M200046-MCP200.

(9) Doherty, M. K.; Hammond, D. E.; Clague, M. J.; Gaskell, S. J.; Beynon, R. J. Turnover of the Human Proteome: Determination of Protein Intracellular Stability by Dynamic SILAC. J. Proteome Res. 2009, 8 (1), 104–112. https://doi.org/10.1021/pr800641v.

(10) Selbach, M.; Schwanhäusser, B.; Thierfelder, N.; Fang, Z.; Khanin, R.; Rajewsky, N. Widespread Changes in Protein Synthesis Induced by MicroRNAs. Nature 2008, 455 (7209), 58–63. https://doi.org/10.1038/nature07228.

(11) Schwanhäusser, B.; Gossen, M.; Dittmar, G.; Selbach, M. Global Analysis of Cellular Protein Translation by Pulsed SILAC. Proteomics 2009, 9 (1), 205–209. https://doi.org/10.1002/pmic.200800275.

(12) Stepath, M.; Zülch, B.; Maghnouj, A.; Schork, K.; Turewicz, M.; Eisenacher, M.; Hahn, S.; Sitek, B.; Bracht, T. Systematic Comparison of Label-Free, SILAC, and TMT Techniques to Study Early Adaption toward Inhibition of EGFR Signaling in the Colorectal Cancer Cell Line DiFi. J. Proteome Res. 2020, 19 (2), 926–937. https://doi.org/10.1021/acs.jproteome.9b00701.

(13) Salovska, B.; Zhu, H.; Gandhi, T.; Frank, M.; Li, W.; Rosenberger, G.; Wu, C.; Germain, P.-L.; Zhou, H.; Hodny, Z.; Reiter, L.; Liu, Y. Isoform-Resolved Correlation Analysis between MRNA Abundance Regulation and Protein Level Degradation. Mol. Syst. Biol. 2020, 16 (3), e9170. https://doi.org/10.15252/msb.20199170.

(14) Pino, L. K.; Baeza, J.; Lauman, R.; Schilling, B.; Garcia, B. A. Improved SILAC Quantification with Data-Independent Acquisition to Investigate Bortezomib-Induced Protein Degradation. J. Proteome Res. 2021, 20 (4), 1918–1927. https://doi.org/10.1021/acs.jproteome.0c00938.

(15) Wu, C.; Ba, Q.; Lu, D.; Li, W.; Salovska, B.; Hou, P.; Mueller, T.; Rosenberger, G.; Gao, E.; Di, Y.; Zhou, H.; Fornasiero, E. F.; Liu, Y. Global and Site-Specific Effect of Phosphorylation on Protein Turnover. Dev. Cell 2021, 56 (1), 111–124.e6. https://doi.org/10.1016/j.devcel.2020.10.025.

(16) Salovska, B.; Li, W.; Di, Y.; Liu, Y. BoxCarmax: A High-Selectivity Data-Independent Acquisition Mass Spectrometry Method for the Analysis of Protein Turnover and Complex Samples. Anal. Chem. 2021, 93 (6), 3103–3111. https://doi.org/10.1021/acs.analchem.0c04293.

(17) Meier, F.; Geyer, P. E.; Virreira Winter, S.; Cox, J.; Mann, M. BoxCar Acquisition Method Enables Single-Shot Proteomics at a Depth of 10,000 Proteins in 100 Minutes. Nat. Methods 2018, 15 (6), 440–448. https://doi.org/10.1038/s41592-018-0003-5.

(18) Egertson, J. D.; Kuehn, A.; Merrihew, G. E.; Bateman, N. W.; MacLean, B. X.; Ting, Y. S.; Canterbury, J. D.; Marsh, D. M.; Kellmann, M.; Zabrouskov, V.; Wu, C. C.; MacCoss, M. J. Multiplexed MS/MS for Improved Data-Independent Acquisition. Nat. Methods 2013, 10 (8), 744–746. https://doi.org/10.1038/nmeth.2528.

(19) Vincent, C. E.; Potts, G. K.; Ulbrich, A.; Westphall, M. S.; Atwood, J. A.; Coon, J. J.; Weatherly, D. B. Segmentation of Precursor Mass Range Using ‘Tiling’ Approach Increases Peptide Identifications for MS1-Based Label-Free Quantification. Anal. Chem. 2013, 85 (5), 2825–2832. https://doi.org/10.1021/ac303352n.

(20) Ludwig, C.; Gillet, L.; Rosenberger, G.; Amon, S.; Collins, B. C.; Aebersold, R. Data-Independent Acquisition-Based SWATH-MS for Quantitative Proteomics: A Tutorial. Mol. Syst. Biol. 2018, 14 (8), e8126. https://doi.org/10.15252/msb.20178126.

(21) Graumann, J.; Scheltema, R. A.; Zhang, Y.; Cox, J.; Mann, M. A Framework for Intelligent Data Acquisition and Real-Time Database Searching for Shotgun Proteomics. Mol. Cell. Proteomics MCP 2012, 11 (3), M111.013185. https://doi.org/10.1074/mcp.M111.013185.

(22) Meyer, J. G.; Niemi, N. M.; Pagliarini, D. J.; Coon, J. J. Quantitative Shotgun Proteome Analysis by Direct Infusion. Nat. Methods 2020, 17 (12), 1222–1228. https://doi.org/10.1038/s41592-020-00999-z.

(23) Shevchenko, A.; Chernushevich, I.; Ens, W.; Standing, K. G.; Thomson, B.; Wilm, M.; Mann, M. Rapid ‘de Novo’ Peptide Sequencing by a Combination of Nanoelectrospray, Isotopic Labeling and a Quadrupole/Time-of-Flight Mass Spectrometer. Rapid Commun. Mass Spectrom. 1997, 11 (9), 1015–1024. https://doi.org/10.1002/(SICI)1097-0231(19970615)11:9<1015::AID-RCM958>3.0.CO;2-H

(24) Qin, J.; Herring, C. J.; Zhang, X. De Novo Peptide Sequencing in an Ion Trap Mass Spectrometer with 18O Labeling. Rapid Commun. Mass Spectrom. 1998, 12 (5), 209–216. https://doi.org/10.1002/(SICI)1097-0231(19980314)12:5<209::AID-RCM141>3.0.CO;2-S.

(25) Goodlett, D. R.; Keller, A.; Watts, J. D.; Newitt, R.; Yi, E. C.; Purvine, S.; Eng, J. K.; Haller, P. von; Aebersold, R.; Kolker, E. Differential Stable Isotope Labeling of Peptides for Quantitation and de Novo Sequence Derivation. Rapid Commun. Mass Spectrom. 2001, 15 (14), 1214–1221. https://doi.org/10.1002/rcm.362.

(26) Volchenboum, S. L.; Kristjansdottir, K.; Wolfgeher, D.; Kron, S. J. Rapid Validation of Mascot Search Results via Stable Isotope Labeling, Pair Picking, and Deconvolution of Fragmentation Patterns*. Mol. Cell. Proteomics 2009, 8 (8), 2011–2022. https://doi.org/10.1074/mcp.M800472-MCP200.

(27) Heller, M.; Mattou, H.; Menzel, C.; Yao, X. Trypsin Catalyzed 16O-to-18O Exchange for Comparative Proteomics: Tandem Mass Spectrometry Comparison Using MALDI-TOF, ESI-QTOF, and ESI-Ion Trap Mass Spectrometers. J. Am. Soc. Mass Spectrom. 2003, 14 (7), 704–718. https://doi.org/10.1016/S1044-0305(03)00207-1.

(28) Bamberger, C.; Pankow, S.; Park, S. K. R.; Yates, J. R. Interference-Free Proteome Quantification with MS/MS-Based Isobaric Isotopologue Detection. J. Proteome Res. 2014, 13 (3), 1494–1501. https://doi.org/10.1021/pr401035z.

(29) Merrill, A. E.; Hebert, A. S.; MacGilvray, M. E.; Rose, C. M.; Bailey, D. J.; Bradley, J. C.; Wood, W. W.; El Masri, M.; Westphall, M. S.; Gasch, A. P.; Coon, J. J. NeuCode Labels for Relative Protein Quantification*. Mol. Cell. Proteomics 2014, 13 (9), 2503–2512. https://doi.org/10.1074/mcp.M114.040287.

(30) Eng, J. K.; Jahan, T. A.; Hoopmann, M. R. Comet: An Open-Source MS/MS Sequence Database Search Tool. Proteomics 2013, 13 (1), 22–24. https://doi.org/10.1002/pmic.201200439.

(31) Eng, J. K.; Hoopmann, M. R.; Jahan, T. A.; Egertson, J. D.; Noble, W. S.; MacCoss, M. J. A Deeper Look into Comet—Implementation and Features. J. Am. Soc. Mass Spectrom. 2015, 26 (11), 1865–1874. https://doi.org/10.1007/s13361-015-1179-x.

(32) Eng, J. K.; Fischer, B.; Grossmann, J.; MacCoss, M. J. A Fast SEQUEST Cross Correlation Algorithm. J. Proteome Res. 2008, 7 (10), 4598–4602. https://doi.org/10.1021/pr800420s.

(33) Chambers, M. C.; Maclean, B.; Burke, R.; Amodei, D.; Ruderman, D. L.; Neumann, S.; Gatto, L.; Fischer, B.; Pratt, B.; Egertson, J.; Hoff, K.; Kessner, D.; Tasman, N.; Shulman, N.; Frewen, B.; Baker, T. A.; Brusniak, M.-Y.; Paulse, C.; Creasy, D.; Flashner, L.; Kani, K.; Moulding, C.; Seymour, S. L.; Nuwaysir, L. M.; Lefebvre, B.; Kuhlmann, F.; Roark, J.; Rainer, P.; Detlev, S.; Hemenway, T.; Huhmer, A.; Langridge, J.; Connolly, B.; Chadick, T.; Holly, K.; Eckels, J.; Deutsch, E. W.; Moritz, R. L.; Katz, J. E.; Agus, D. B.; MacCoss, M.; Tabb, D. L.; Mallick, P. A Cross-Platform Toolkit for Mass Spectrometry and Proteomics. Nat. Biotechnol. 2012, 30 (10), 918–920. https://doi.org/10.1038/nbt.2377.

(34) Elias, J. E.; Gygi, S. P. Target-Decoy Search Strategy for Increased Confidence in Large-Scale Protein Identifications by Mass Spectrometry. Nat. Methods 2007, 4 (3), 207–214. https://doi.org/10.1038/nmeth1019.

(35) Teleman, J.; Chawade, A.; Sandin, M.; Levander, F.; Malmström, J. Dinosaur: A Refined Open-Source Peptide MS Feature Detector. J. Proteome Res. 2016, 15 (7), 2143–2151. https://doi.org/10.1021/acs.jproteome.6b00016.

(36) Goloborodko, A. A.; Levitsky, L. I.; Ivanov, M. V.; Gorshkov, M. V. Pyteomics—a Python Framework for Exploratory Data Analysis and Rapid Software Prototyping in Proteomics. J. Am. Soc. Mass Spectrom. 2013, 24 (2), 301–304. https://doi.org/10.1007/s13361-012-0516-6.

(37) Levitsky, L. I.; Klein, J. A.; Ivanov, M. V.; Gorshkov, M. V. Pyteomics 4.0: Five Years of Development of a Python Proteomics Framework. J. Proteome Res. 2019, 18 (2), 709–714. https://doi.org/10.1021/acs.jproteome.8b00717.

(38) Fondrie, W. E.; Noble, W. S. Mokapot: Fast and Flexible Semisupervised Learning for Peptide Detection. J. Proteome Res. 2021, 20 (4), 1966–1971. https://doi.org/10.1021/acs.jproteome.0c01010.

(39) Bakalarski, C. E.; Elias, J. E.; Villén, J.; Haas, W.; Gerber, S. A.; Everley, P. A.; Gygi, S.P. The Impact of Peptide Abundance and Dynamic Range on Stable-Isotope-Based Quantitative Proteomic Analyses. J. Proteome Res. 2008, 7 (11), 4756–4765. https://doi.org/10.1021/pr800333e.

(40) Yergey, James.; Heller, David.; Hansen, Gordon.; Cotter, R. J.; Fenselau, Catherine. Isotopic Distributions in Mass Spectra of Large Molecules. Anal. Chem. 1983, 55 (2), 353–356. https://doi.org/10.1021/ac00253a037.

(41) Chavez, J. D.; Keller, A.; Mohr, J. P.; Bruce, J. E. Isobaric Quantitative Protein Interaction Reporter Technology for Comparative Interactome Studies. Anal. Chem. 2020, 92 (20), 14094–14102. https://doi.org/10.1021/acs.analchem.0c03128.

(42) Bittremieux, W. Spectrum_utils: A Python Package for Mass Spectrometry Data Processing and Visualization. Anal. Chem. 2020, 92 (1), 659–661. https://doi.org/10.1021/acs.analchem.9b04884.

(43) Venable, J. D.; Dong, M.-Q.; Wohlschlegel, J.; Dillin, A.; Yates, J. R. Automated Approach for Quantitative Analysis of Complex Peptide Mixtures from Tandem Mass Spectra. Nat. Methods 2004, 1 (1), 39–45. https://doi.org/10.1038/nmeth705.

(44) Haynes, S. E.; Majmudar, J. D.; Martin, B. R. DIA-SIFT: A Precursor and Product Ion Filter for Accurate Stable Isotope Data-Independent Acquisition Proteomics. Anal. Chem. 2018, 90 (15), 8722–8726. https://doi.org/10.1021/acs.analchem.8b01618.

(45) Schweppe, D. K.; Eng, J. K.; Yu, Q.; Bailey, D.; Rad, R.; Navarrete-Perea, J.; Huttlin, E. L.; Erickson, B. K.; Paulo, J. A.; Gygi, S. P. Full-Featured, Real-Time Database Searching Platform Enables Fast and Accurate Multiplexed Quantitative Proteomics. J. Proteome Res. 2020, 19 (5), 2026–2034. https://doi.org/10.1021/acs.jproteome.9b00860.

