## Supplementary Material for "Coisolation of peptide pairs for peptide identification and MS/MS-based quantification"

### SUPPLEMENTARY DISCUSSION

***Consideration of b-ion fragments for Coiso SILAC Comet database search***

Unique to coisolation searches, missed cleavages can result in not only paired y-ions, but also paired b-ions. Particularly of note, when performing a Comet internal decoy search, candidate peptides ending in two lysines will generate decoys when the sequence is pseudo reversed with double the *in silico* b-ion fragments possible for calculating an Xcorr, thus providing a bias to missed cleaved decoys and additionally all peptide candidates that have a missed cleavage.

For generating a coisolation search algorithm, we tested generating theoretical SILAC paired spectra for peptide spectral matching that contained paired y-ions and paired b-ions if possible (silac_pair_fragments=1) and a strategy that only considers paired y-ions and only light or heavy b-ions (for light and heavy triggered peptides respectively; silac_pair_fragments=2). To test this hypothesis, we generated a 1-to-1 SILAC (Lys0:Lys8) *S. cerevisiae* proteome mixture that was digested with Lys-C protease. Using our modified DDA coisolation schema, we triggered three scans on the same precursor: isolated 4 Da offset to the left when heavy (1) and to the right when light (2) with 6.5 m/z wide window scans, and our DDA control being an 1.6 m/z narrow window scan (3) targeting the single light or heavy precursor (**Fig 1a**), giving the resultant MS/MS spectra (**Fig 1b**).

The correctly coisolated wide window scan (left or right) was coisolation searched with Comet considering (silac_pair_fragments=1) or excluding (silac_pair_fragments=2) possible paired b-ions. Both coisolation searches were compared to the narrow window scan searched spectra with the traditional non-paired DDA Comet parameters for each targeted precursor. The proportion of decoy SILAC pairs that did not have exactly one lysine were over-represented in the wide window-coisolation search considering b-ion pairs as compared to the similar proportion observed in wide window-coisolation search excluding b-ion pairs and narrow window-DDA search (**Supp. Fig 1a**). This observation suggests a bias toward missed cleaved candidate peptides (target or decoy) when including paired b-ion fragments. Additionally, the missed cleavage PSM bias is reflected in lower identifications at a PSM 1% FDR based on E-values (**Supp. Fig 1b**).

Of note, E-values are strongly influenced by the tail of the cumulative distribution function of the candidate Xcorr distribution for each scan. Considering the missed cleavage b-ion bias will likely influence a subset of candidates towards this tail broadening the Xcorr distribution, one could expect that the coisolation search including possible paired b-ions will decrease target PSMs $-\log_{10}$(E-values) (higher is better). This notion was supported by the observation that the E-values from the coisolation search excluding paired b-ion outperformed the coisolation search including possible b-ion pairs greater than 2-fold of the time comparing the same spectra with the same matched PSM (**Supp. Fig 1c**). Therefore, we utilized the coisolation search algorithm that excludes the possible paired b-ions for it standardizes the possible theoretical fragments based on peptide length across all possible candidates, generates more PSMs, and improves E-values.

### SUPPLEMENTARY FIGURES

#### *Supplementary Figure 1: Inclusion or exclusion of paired b-ions for peptide-spectral matching*

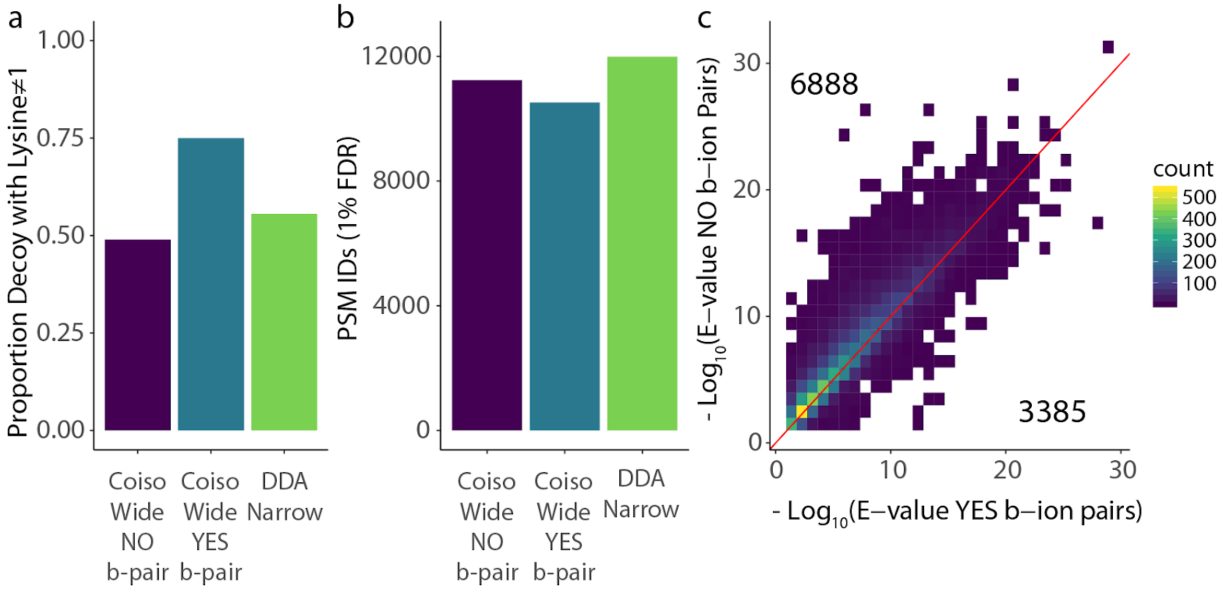


**Supplementary Figure 1: Coiso SILAC Comet searches excluding b-ion pairs removes search bias. a)** Analysis from a *S. cerevisiae* 1:1 SILAC ratio proteome mixture. Bar plot of the proportion of decoy hits that have lysine not equal to 1. The combinations include Coiso Wide searches including or excluding b-ion paired fragments and the traditional DDA search with narrow isolation, all considering the same targeted precursor m/z. **b)** Bar plot of PSMs at 1% FDR based on Comet E-value for the same acquisition search combinations as in (**a**). **c)** Density-binned scatter plot comparing Comet standard E-value including (x-axis) and excluding (y-axis) b-ion paired fragments for matching PSM identification and triggered precursor m/z (filtered in both searches at 1% FDR based on E-value).

### ***Supplementary Figure 2: E-values for DDA and Coiso SILAC analysis***

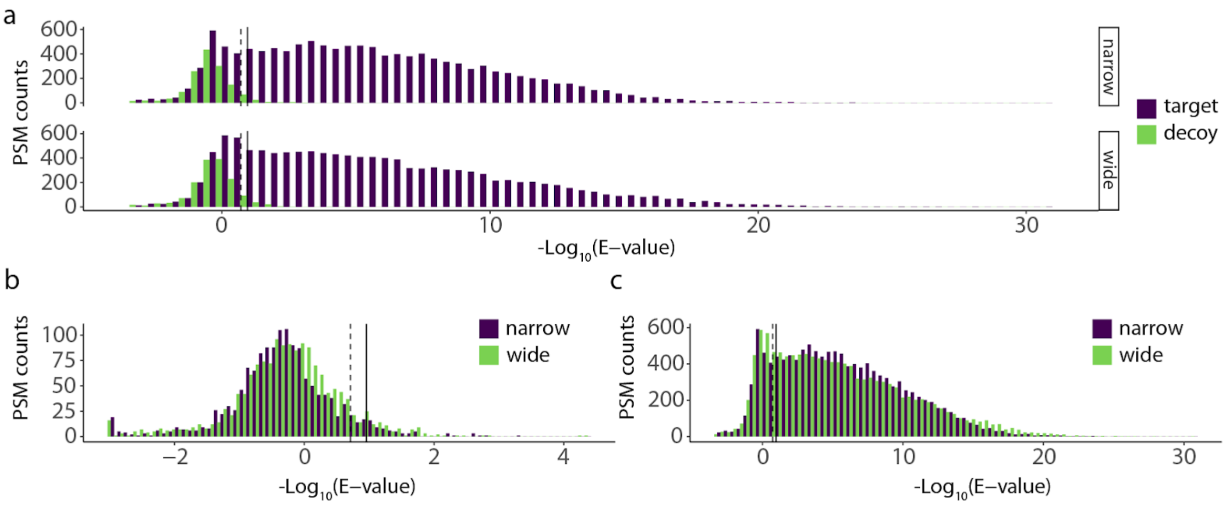


**Supplementary Figure 2: Coiso SILAC E-values shifted for targets and decoys. a)** For *S. cerevisiae* 1:1 SILAC ratio proteome mixture, histograms for PSM counts for targets (purple) and decoys (green) for narrow window scans searched with traditional DDA Comet parameters (top) and for wide window offset scans searched with Coiso Comet parameters (bottom). E-value 1% PSM FDR cut-offs are designated by vertical lines (dashed : DDA Narrow; solid: Coiso Wide). **b)** Histograms for PSM decoys of (**a**) for both MS acquisition-search combinations overlaid. Wide window scans with Coiso search in green and narrow window scans with traditional DDA search in purple, with respective 1% PSM FDR designations as in (**a**). **c)** Histograms for PSM targets as in (**a**) for both MS acquisition-search combinations overlaid. Same color designations as in (**b**) and 1% PSM FDR cut-offs as in (**a**).

### ***Supplementary Figure 3: Coiso E-value calculation***

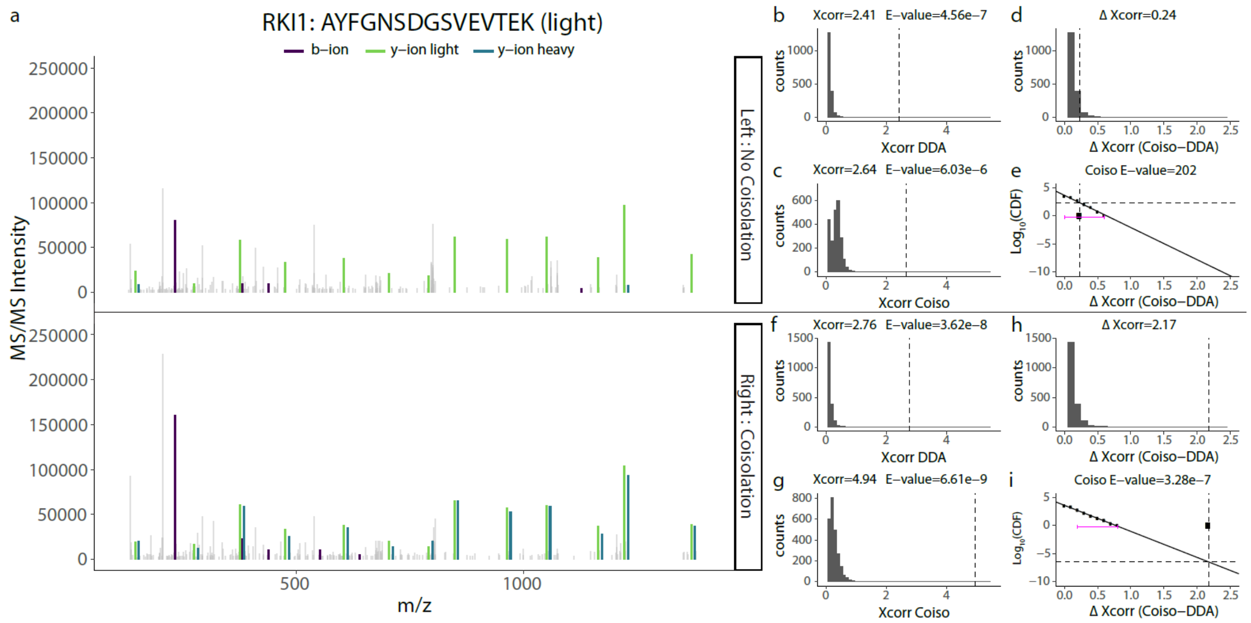


**Supplementary Figure 3: Coiso E-value calculation with a left and right Coiso scan example. a)** Left (top panel) and right (bottom panel) offset wide window Coiso scans for the triggered RKI1 light peptide (zoomed in over fragment ions; dropping dominant non-PSM peaks) with annotated PSM b-ions (purple), light y-ions (green), and heavy y-ions (blue). We can visualize the y-ion Xcorr from the non-targeted precursor by calculating the difference between Coiso Xcorr (precursor pair y-ions and b-ions) minus DDA Xcorr (targeted precursor y-ions and b-ions). The difference results in the delta Xcorr (Coiso - DDA) being the Xcorr contribution from the non-targeted precursor y-ion Xcorr. **b)** Histogram of PSM candidates' Comet Xcorrs when searched with traditional DDA search parameters (only light b- and y-ions) for left offset Coiso scan (from **a**: top panel) with dashed vertical line designating the Xcorr for the PSM sequence identification in (**a**). PSM's Xcorr and E-value for DDA search are noted. **c)** Same as (**b**) for the same PSM candidates when spectra are searched with Coiso SILAC Comet parameters. PSM's Xcorr and E-value for Cosio search are noted. **d)** Same as (**b**) for the same PSM candidates for delta Xcorr (Coiso Xcorr - DDA Xcorr). Coiso Xcorr from (**c**) minus DDA Xcorr from (**b**) for matching candidates. PSM's delta Xcorr is noted. **e)** Scatter plot of the logarithmic transform of the cumulative distribution function (CDF) based on the histogram in (**c**) binned by 0.1 delta Xcorr. Linear regression trendline (solid line) fit on the right tail of delta Xcorr $\log_{10}$(CDF) with the range defined by magenta brackets for E-value calculation. The horizontal dashed line defines the point of intersection between the PSM's delta Xcorr (vertical dashed line and point on the x-axis) and the linear regression trendline which is used to extrapolate the Coiso E-value noted on the plot. **f-i)** Same as (**b-e**) for the right offset Coiso scan.

### *Supplementary Figure 4: E-value comparing Coiso scans between offsets*

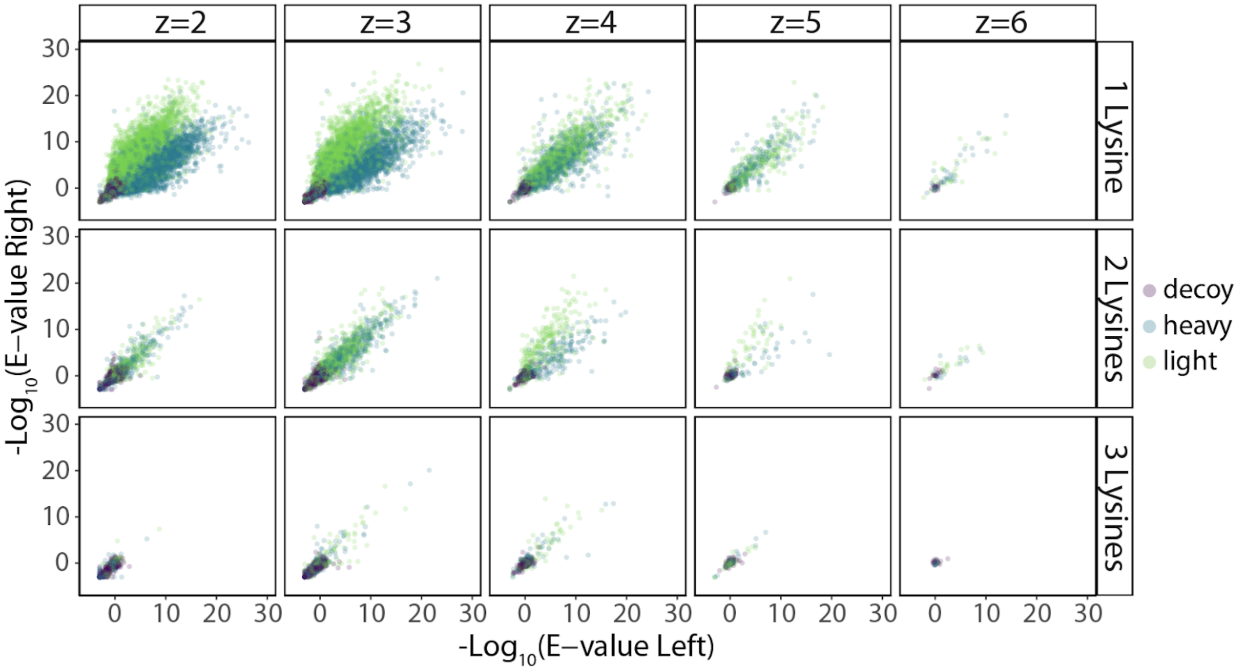


**Supplementary Figure 4: Comet E-value can marginally distinguish between successful and unsuccessful SILAC peptide pair coisolation.** Scatterplot for standard Comet E-value (from Coiso search parameters) of the 1:1 *S. cerevisiae* SILAC proteome mixture faceted by PSM charge state and number of lysines for matching left and right offset Coiso scans. PSM assignment either heavy (blue), light (green), or decoy (purple) based on correctly Coiso scans PSM sequence.

### *Supplementary Figure 5: Number of quantifiable MS1 vs MS/MS features*

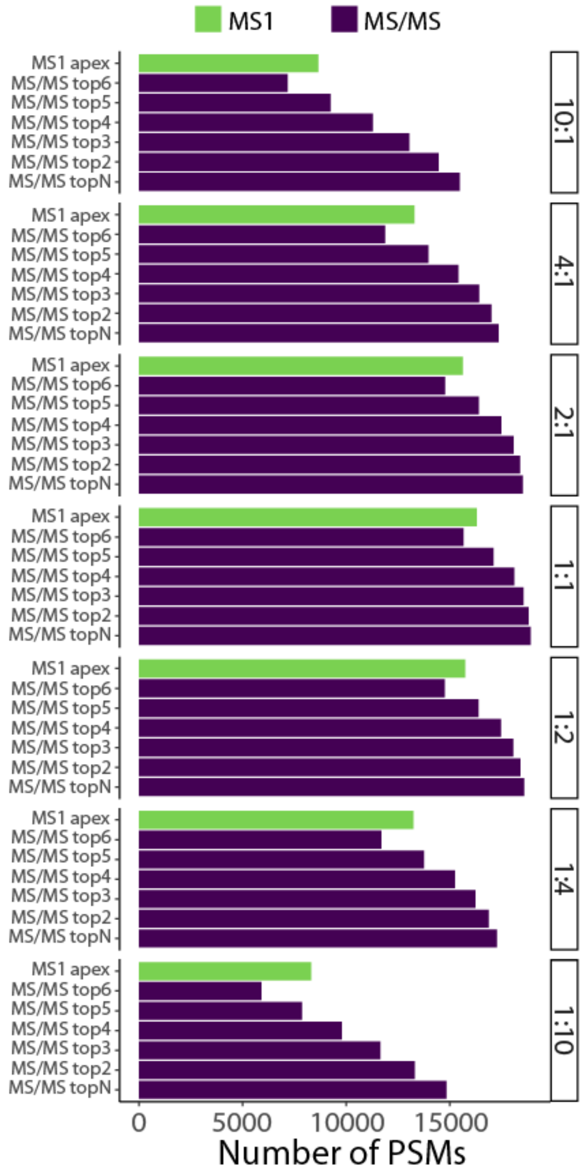


**Supplementary Figure 5: Coiso SILAC MS/MS captures more quantifiable peptide-spectral matches than MS1.** Bar plot of quantifiable peptide-spectral matches for summed or apex-based MS1 features (green) from Dinosaur or Coiso MS/MS-based quantifications (purple) for top2-top6 and topN (all matched) fragment ions. Plots are annotated with the Lys0:Lys8 SILAC ratio (Lys0:Lys8 respectively).

### *Supplementary Figure 6: Overlap of quantifiable MS1 and Coiso MS/MS features*

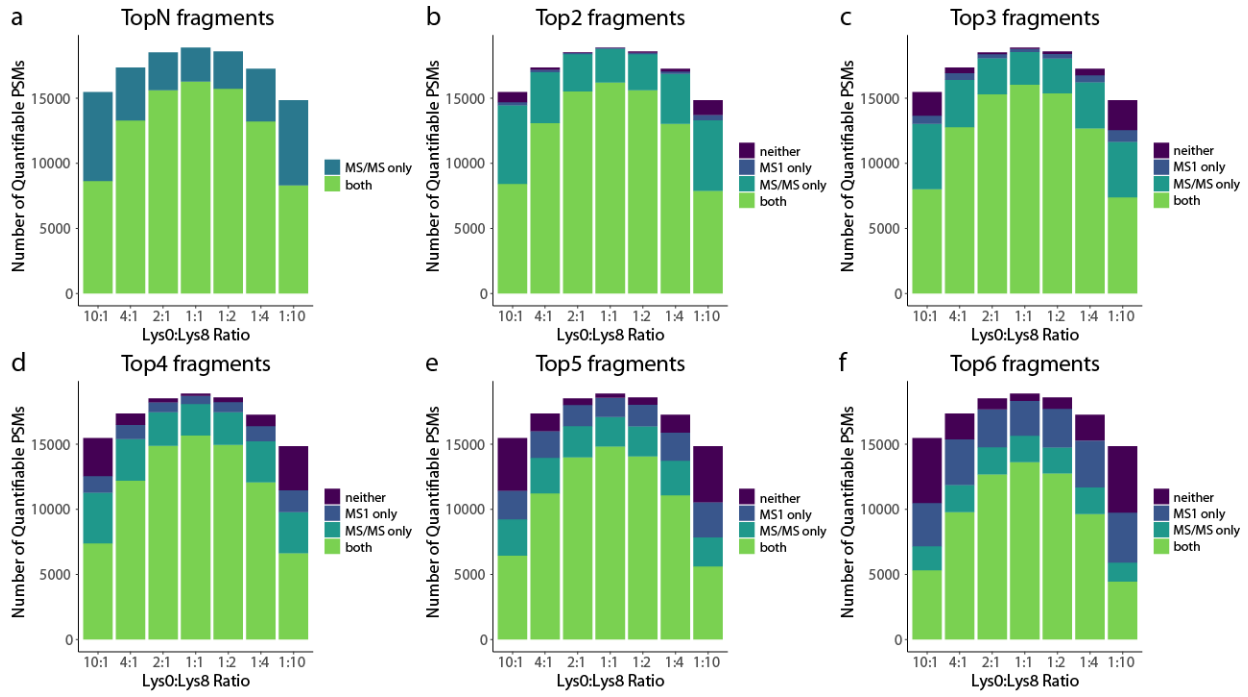


**Supplementary Figure 6: Coiso SILAC method overlap with MS1 quantifications varies based on number of quantifiable fragment filters. (a-f)** Stacked bar plots of the number of quantifiable PSMs colored by overlap designation (both: quantifiable MS1 and MS/MS (green); neither: not quantifiable in MS1 and MS/MS (purple); MS1 only (dark blue); MS/MS only (light blue)). Each panel refers to the number of quantifiable fragments required to ensure an MS/MS quantifiable PSM (**a**:topN; **b**:top2; **c**:top3; **d**:top4; **e**:top5; **f**:top6). Plots are generated based on SILAC-labeled *S. cerevisiae* proteome mixtures with defined SILAC ratios of Lys0:Lys8 respectively.

### *Supplementary Figure 7: MS1 and MS/MS SILAC ratio quantification distributions*

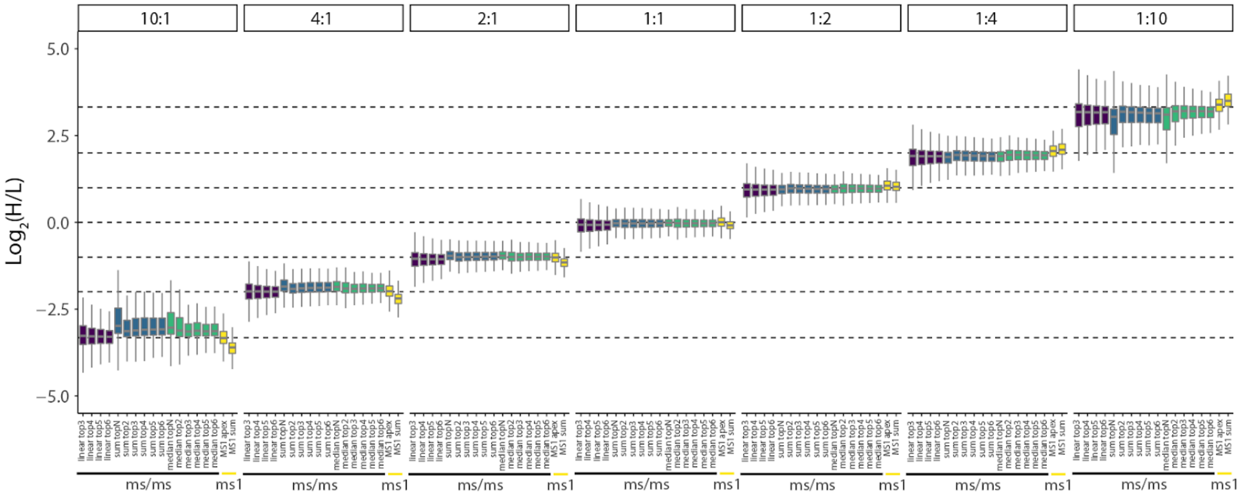


**Supplementary Figure 7: Coiso SILAC quantification methods and filters vary in precision and accuracy.** Box plots for peptide-spectral matches across SILAC *S.cerevisiae* proteome mixtures (Lys0:Lys8 respectively). Three MS/MS-based quantification methods were used: median-based (green), sum-based (blue), and linear regression-based (purple) filtered based on the topN or top2-6 quantifiable paired y-ion fragments. MS1-matched SILAC features from Dinosaur (yellow) using either apex or sum-based quantification. Box plots represent the distribution from all quantifiable PSMs from Supplementary Figure 6. In the box plot, the horizontal line represents the median, box designates the IQR, and the whiskers indicate 1.5 x IQR from the box ends.

### *Supplementary Figure 8: MS1 vs MS/MS quantifications across dynamic range*

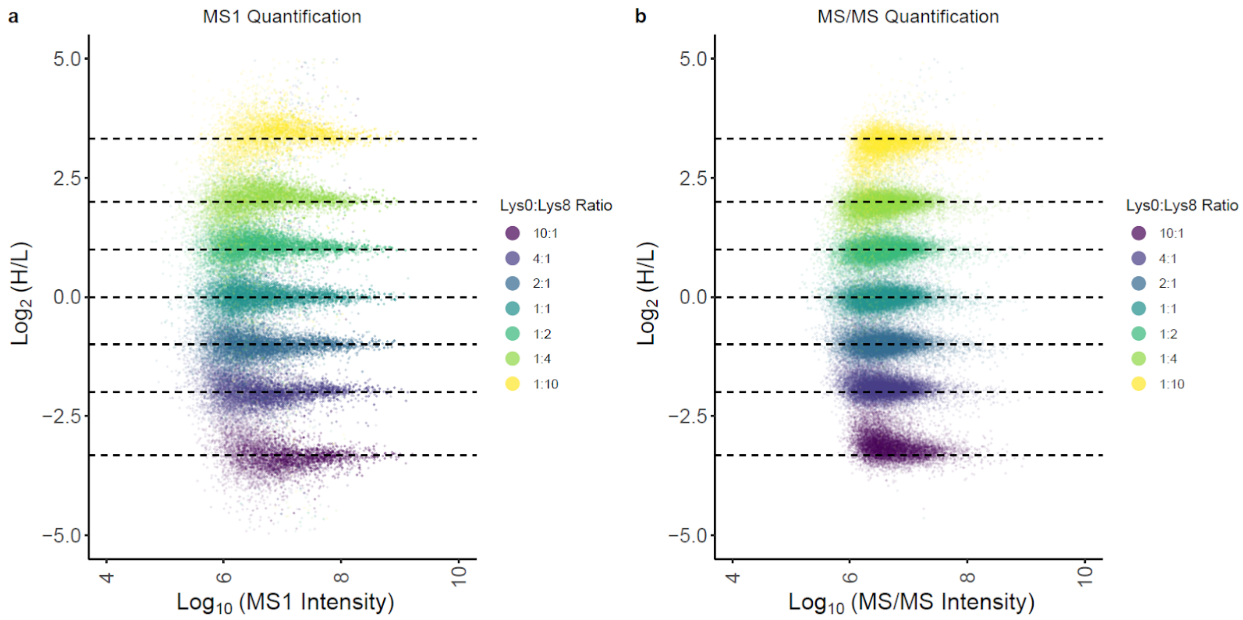


**Supplementary Figure 8: Coiso SILAC MS/MS and MS1 quantifications map to expected ratios across dynamic range.** Scatter plot of all quantifiable PSMs colored by SILAC sample for SILAC *S. cerevisiae* proteome mixtures (Lys0:Lys8 ratio respectively). MS1 intensities are calculated via the sum of heavy and light peptide feature's intensities. MS/MS intensities are derived from the sum of all the peptides' (heavy and light) fragment ion signals from the single MS/MS of the peptide-spectral match.

### *Supplementary Figure 9: Percentage of outliers for all MS1 and MS/MS quantification strategies*

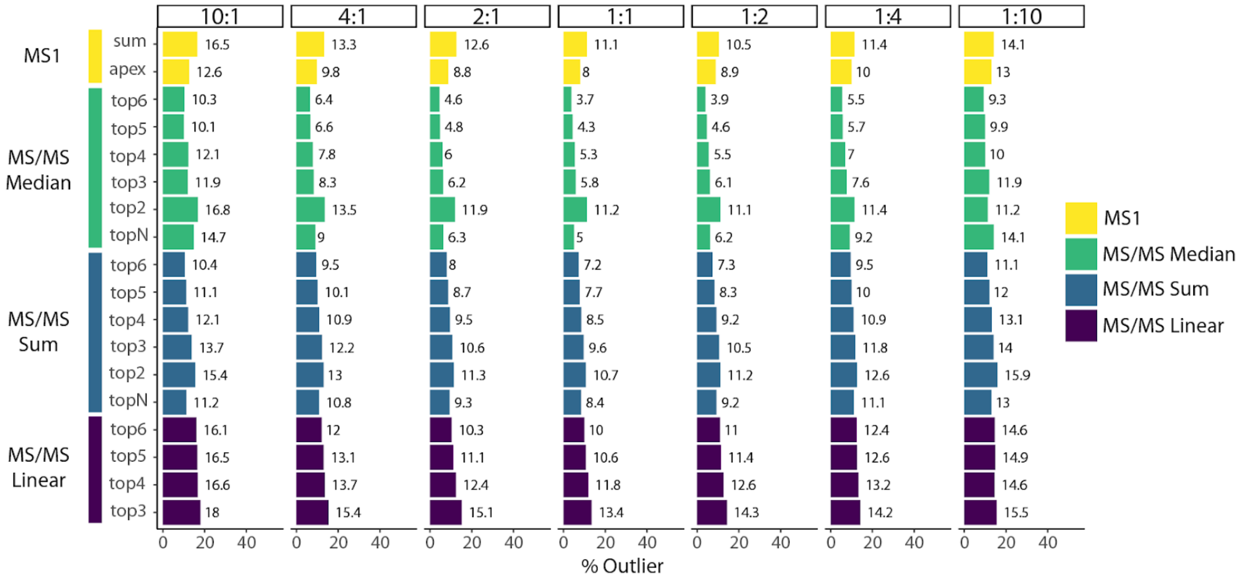


**Supplementary Figure 9: Coiso SILAC quantification methods and filters vary in outliers.** Bar plots of percentage peptide-spectral matches that are outliers from the SILAC *S.cerevisiae* proteome mixture distribution (Lys0:Lys8 respectively). MS/MS quantification methods (median-based:green, sum-based:blue, linear-based:purple) filtered based on the number of quantifiable paired y-ion fragments and Dinosaur MS1-derived quantifications (yellow) are represented here. Outliers are PSM $log_2$(ratios) > Q3 + (1.5 x IQR) or $log_2$(ratios) < Q1 - (1.5 x IQR).

### ***Supplementary Figure 10: Comparing Lys0:Lys8 and Lys6:Lys8 SILAC mixture's quantifiable identifications and quantification precision and accuracy.***

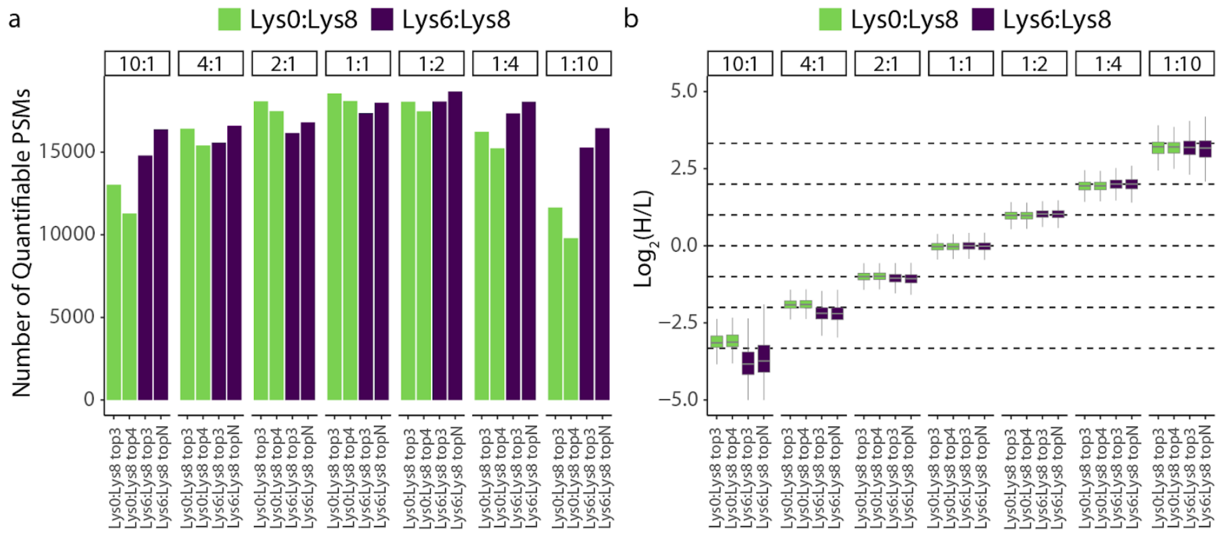


**Supplementary Figure 10: Lys0:Lys8 vs. Lys6:Lys8 Coiso SILAC MS/MS quantification comparison of quantifiable PSMs and quantification distributions. a)** Bar plot of quantifiable peptide-spectral matches for SILAC-labeled *S. cerevisiae* Lys0:Lys8 (green) and Lys6:Lys8 (purple) proteome mixtures using Coiso MS/MS-based qualifications with respective fragment ion filters. Plots are faceted by sample with defined SILAC ratio (matching Lys0:Lys8 and Lys6:Lys8 respectively) of a SILAC-labeled *S. cerevisiae* proteome mixture. **b)** Box plots of SILAC ratio for the distribution of all quantifiable PSMs from(**a**). MS/MS-based quantification methods from Lys0:Lys8 (green) and Lys6:Lys8 (purple) filtered by respective quantifiable paired y-ion fragments. In the box plot, the horizontal line represents the median, box designates the IQR, and the whiskers indicate 1.5 x IQR from the box ends.

*Supplementary Figure 11: Lys6:Lys8 deconvolution algorithm overcorrects light contribution in 10:1 sample when theoretical heavy contribution is smaller*

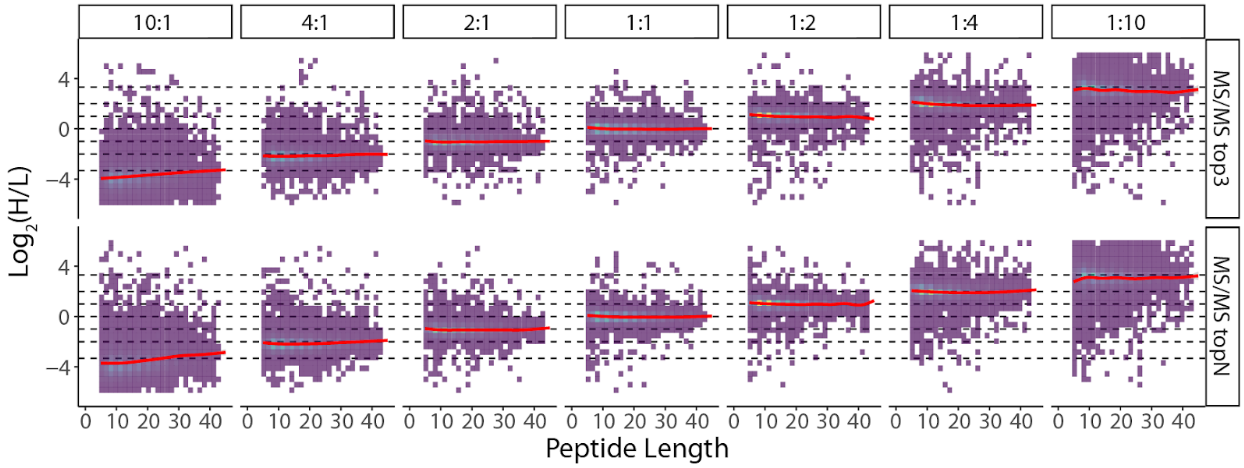


**Supplementary Figure 11: Lys6:Lys8 Coiso SILAC MS/MS quantification changes associated with peptide size for the 10:1 Lys6:Lys8 sample.** Density-binned scatter plot of Coiso SILAC MS/MS deconvolution quantifications across peptide length for median top3 and topN fragment ion filters. Dashed lines designated expected theoretical SILAC ratios. Red trendline represents the data fit to a general additive model (GAM).
